# Visual deprivation independent shift of ocular dominance induced by cross-modal plasticity

**DOI:** 10.1101/413229

**Authors:** Manuel Teichert, Marcel Isstas, Lutz Liebmann, Christian A. Hübner, Franziska Wieske, Christine Winter, Jürgen Bolz

## Abstract

There is convincing evidence that the deprivation of one sense can lead to adaptive neuronal changes in the spared primary sensory cortices. However, the repercussions of late-onset sensory deprivations on functionality of the remaining sensory cortices are poorly understood. Using repeated intrinsic signal imaging we investigated the effects of whisker or auditory deprivation (WD or AD, respectively) on responsiveness of the binocular primary visual cortex (V1) in fully adult mice. The binocular zone of mice is innervated by both eyes, with the contralateral eye always dominating V1 input over ipsilateral eye input, the normal ocular dominance (OD) ratio. Strikingly, we found that 3 days after WD or AD there was a transient shift of OD, which was mediated by a potentiation of V1 input through the ipsilateral eye. This cross-modal effect was accompanied by strengthening of V1 layer 4 synapses, required visual experience through the ipsilateral eye and was mediated by an increase of the excitation/inhibition ratio in V1. Finally, we demonstrate that both WD and AD induced a long-lasting improvement of visual performance. Our data provide evidence that the deprivation of a non-visual sensory modality cross-modally induces experience dependent V1 plasticity and improves visual behavior, even in adult mice.

## Introduction

It has been demonstrated that the loss or deprivation of one sensory modality can have striking effects on the remaining senses. Such changes are broadly referred as to “cross-modal plasticity” and can improve the functionality of the spared senses (Neville & Lawson, 1987; Cohen *et al.*, 1997; Lomber *et al.*, 2010; Lee & Whitt, 2015; Teichert & Bolz, 2017b; Teichert *et al.*, 2018a). It has been suggested that these compensatory enhancements arise because the deprived cortex becomes driven by the spared sensory modalities (Cohen *et al.*, 1997; Lomber *et al.*, 2010).

However, there is increasing evidence that functional improvements of the spared senses can be attributed to adaptive changes in the spared sensory cortices. For instance, we could recently demonstrate that auditory deprivation (AD) leads to a rapid increase of visually evoked responses in the spared V1 which was accompanied by improvements of V1 spatial frequency and contrast tuning (Teichert & Bolz, 2017b; a). While these changes appeared most likely due to a rapid disinhibitory effect (Teichert & Bolz, 2017b), previous studies could demonstrate that more prolonged sensory deprivations lead to plastic alterations in the spared primary sensory cortices (Lee & Whitt, 2015). For example, 2 days of visual deprivation in juvenile mice selectively strengthened layer 4-2/3 synapses in the somatosensory barrel cortex and sharpened the functional whisker-barrel map at layer 2/3 (Jitsuki *et al.*, 2011). Similarly, 7 days of visual deprivation was shown to strengthen thalamo-cortical synapses in the spared primary auditory cortex (A1) of juvenile and adult mice (Petrus *et al.*, 2014). These plastic changes were accompanied by increased contrast sensitivity and frequency tuning of A1 neurons (Petrus *et al.*, 2014). Moreover, in a recent study we could demonstrate that 7-14 days of whisker deprivation (WD) in fully adult mice led to massive enhancements of spatial frequency and contrast tuning of the primary visual cortex (V1), which were markedly stronger compared to the enhancements observed after acute sensory deprivations (Teichert & Bolz, 2017b; Teichert *et al.*, 2018a). We could further show that both WD and AD act to restore ocular dominance (OD) plasticity in the spared V1 in fully adult mice older than 120 days (Teichert *et al.*, 2018b). Specifically, combing WD or AD with monocular deprivation (MD) for 7 days induced a shift of the OD in the binocular zone of V1 which was mediated by an increased V1 responsiveness to open eye stimulation (Teichert *et al.*, 2018b). Taken together, these findings suggest that sensory deprivations lead to short-term and, in particular, long-term compensatory neuronal alterations in the spared primary sensory cortices. However, the time course of events taking place in the spared cortices is largely unknown. Moreover, the repercussions of late-onset sensory deprivations on functionality of the remaining sensory cortices are poorly understood.

In order to address these issues we here investigated the cross-modal effects of WD or AD on V1 responsiveness and visually mediated behavior at 0, 3 and 7 days after the deprivation in fully adult mice far beyond their sensory critical periods. For this, we chronically measured V1 responses evoked by visual stimulation of the contralateral and ipsilateral eye using Fourier based periodic intrinsic signal imaging. Notably, we found that both WD and AD induced a marked shift of the OD in V1 after 3 days, which was mediated by a strong increase of V1 responses evoked by visual stimulation of the ipsilateral eye. However, after 7 days of WD or AD V1 input through the ipsilateral eye and thus, the OD completely recovered to baseline levels. Finally, we investigated the effects of WD or AD on behavioral visual performance. We found that both spatial frequency and contrast sensitivity thresholds of the optokinetic reflex (OKR) improved in a V1 dependent and V1 independent manner. Taken together, our results suggest that the deprivation of a non-visual sensory modality induces plastic changes in the binocular zone of V1 and leads to long-lasting improvements of visual performance.

## Material and Methods

### Animals

C57BL/6J (Jackson labs) mice were raised in a group of 2-3 in transparent standard cages (16.5×22.5 cm) on a 12 h light/dark cycle, with food and water available *ad libitum*. Between the chronic experiments each animal was housed alone in a standard cage. The environment in the cage was minimally enriched with cotton rolls and nest material. In our mouse facility the light intensity was about 150-170 lux. As demonstrated recently, these rearing conditions are not sufficient to extend OD plasticity into adulthood (Teichert *et al.*, 2018b). Animal housing in our institution is regularly supervised by veterinaries from the state of Thuringia, Germany. For the present study we used fully adult male and female mice (PD 120-240). All experimental procedures have been performed according to the German Law on the Protection of Animals and the corresponding European Communities Council Directive 2010 (2010/63/EU), and were approved by the Thüringer Landesamt für Lebensmittelsicherheit und Verbraucherschutz (Thuringia State Office for Food Safety and Consumer Protection) under the registration numbers 02-032/16 and 02-050/14. Every effort was made to minimize the number of animals used and their suffering.

### Optical imaging of intrinsic signals

#### Mouse preparation for optical imaging

To investigate the effects of WD (n=32) or AD (n=6) on visually evoked activity of V1 we used Fourier based periodic intrinsic signal imaging (Isstas *et al.*, 2017; Teichert & Bolz, 2017b). In addition, imaging experiments were performed in n=9 untreated control mice. As described previously (Teichert & Bolz, 2017b; Teichert *et al.*, 2018a), animals were initially anesthetized with 4% isoflurane in a mixture of 1:1 O_2_/N_2_O and placed on a heating blanket for maintaining body temperature (37.5°C). Subsequently, mice received injections of chlorprothixene (20 µg/mouse i.m.) and carprofene (5 mg/kg, s.c.). The animal was fixed in a stereotaxic frame and we removed the skin of the left hemisphere to expose the visual cortex. The exposed area was covered with 2.5% agarose in saline and sealed with a standard microscope glass coverslip. Cortical responses were always recorded through the intact skull. During the experiment isoflurane inhalation anesthesia was applied through a plastic mask and maintained at 0.5-0.6%.

#### Mouse preparation for repeated imaging experiments

Repeated intrinsic imaging in the same mice was performed as previously described (Kaneko *et al.*, 2008; Teichert *et al.*, 2018b). Briefly, after the first imaging session the skin was re-sutured and animals were returned to their standard cages. During the subsequent days animals received a daily injection of carprofen (5 mg/kg, s.c.) for pain prevention. Before the next imaging session (day 3 and 7) the skin was re-opened and imaging was performed as described above.

#### Imaging of visual cortex

Responses of mouse primary visual cortex were recorded described previously (Kalatsky & Stryker, 2003; Teichert & Bolz, 2017b; Teichert *et al.*, 2018b). Briefly, the method uses a periodic stimulus that is presented to the animal for some time and cortical responses are extracted by Fourier analysis. In our case, the visual stimulus was a drifting horizontal light bar of 2° width, 100% (or 10%, respectively) contrast and with a temporal frequency of 0.125 Hz. The stimulus was presented on a high refresh rate monitor (Hitachi Accuvue HM 4921-D) placed 25 cm in front of the animal. Visual stimulation was adjusted so that it only appeared in the binocular visual field of the recorded hemisphere (−5° to +15° azimuth, −17° to +60° elevation). The stimulus was presented to the contra or ipsilateral eye or to both eyes for 5 min. Thus, the stimulus was repeated for about 35 times during one presentation period.

#### CCD camera recording procedure

Using a Dalsa 1M30 CCD camera (Dalsa, Waterloo, Canada) with a 135×50 mm tandem lens (Nikon, Inc., Melville, NY), we first recorded images of the surface vascular pattern via illumination with green light (550±2 nm) and, after focusing 600 µm below the pial surface, intrinsic signals were obtained via illumination with red light (610±2 nm). Frames were acquired at a rate of 30 Hz and temporally averaged to 7.5 Hz. The 1024×1024 pixel images were spatially averaged to a 512×512 resolution. We always imaged the left hemisphere of the animals.

#### Data analysis

From the recorded frames the signal was extracted by Fourier analysis at the stimulation frequency and converted into amplitude and phase maps using custom software (Kalatsky & Stryker, 2003). In detail, from a pair of the upward and downward maps, a map with absolute retinotopy and an average magnitude map were computed. For data analysis we used the MATLAB standard as described previously (Cang *et al.*, 2005; Lehmann & Lowel, 2008). The magnitude component represents the activation intensity of the visual cortex. Since high levels of neuronal activity decrease oxygen levels supplied by hemoglobin and since deoxyhemoglobin absorbs more red light (610±2 nm), the reflected light intensity decreases in active cortical regions. Because the reflectance changes are very small (less than 0.1%) all amplitudes are multiplied with 10^4^ so that they can be presented as small positive numbers. Thus, the obtained values are dimensionless. Amplitude maps were obtained by averaging the response amplitudes of individual pixels from maps to upward and downward moving bars. The ocular dominance index was computed as (C-I)/(C+I) with C and I representing the peak response amplitudes of V1 elicited by contralateral eye and ipsilateral eye stimulation, as described previously (Cang *et al.*, 2005; Kaneko *et al.*, 2008). To each condition we took at least three magnitudes of V1 responsiveness and averaged them for data presentation.

### Whisker deprivation (WD) and auditory deprivation (AD)

WD and AD were always performed immediately before the first imaging session or optometry experiments (day 0). WD was performed as described previously (He *et al.*, 2012; Teichert *et al.*, 2018a; Teichert *et al.*, 2018b). Briefly, animals were deeply anesthetized with 2% isoflurane in a mixture of 1:1 O_2_/N_2_O applied through a plastic mask. The eyes of the animal were protected with silicon oil. Whiskers (macro vibrissae) were plucked bilaterally using fine forceps. Subsequently mice received an injection of carprofene (4 mg/kg, s.c.) for pain prevention and were returned to their standard cages. Over the following days whiskers were re-shaved every other day, and animals received a daily administration of carprofene (4 mg/kg, s.c.).

AD was always induced by bilateral malleus removal as described previously (Teichert & Bolz, 2017a; b; Teichert *et al.*, 2018a). Briefly, animals were deeply anesthetized with 2% isoflurane in a mixture of 1:1 O_2_/N_2_O applied through a plastic mask. Additionally, mice received a subcutaneous injection of carprofene (4 mg/kg, s.c.) for pain prevention. The eyes of the animal were protected with silicon oil. The tympanic membrane was punctured and the malleus was removed under visual control through this opening using fine sterilized forceps. Great care was taken to avoid any destruction of the stapes and the oval window which is visible through the hearing canal (see (Tucci *et al.*, 1999)). Over the following days animals received a daily administration of carprofene (4 mg/kg, s.c.).

### Monocular deprivation (MD)

We examined whether the cross-modally induced V1 activity changes depend on patterned visual input. For this, in one group of mice (n=4) we sutured the contra and in another group we sutured the ipsilateral eye (n=5). MD was always performed after the first imaging session, thus, during the same anesthesia like WD. For this, we increased the isoflurane concentration to 2% in a mixture of 1:1 N_2_O and O_2_. Lid margins of the contra or ipsilateral eye, respectively, were trimmed and an antibiotic ointment was applied. Subsequently the eye was sutured. After MD animals received one injection of carprofene (4 mg/kg, s.c.) and were returned to their standard cages. All animals were checked daily to ensure that the sutured eye remained closed during the MD time. Over the following 3 days animals received a daily administration of carprofene (4 mg/kg, s.c.) for pain prevention.

### Electrophysiology

#### Slice preparation for electrophysiological recordings

350-µm-thick brain slices were prepared from 3-month-old mice (control: n=4, 3 d WD: n=3) in preparation aCSF (in mM): 2.5 KCl, 6 MgSO_4_, 1.25 NaH_2_PO_4_, 0.25 CaCl_2_, 260 D-glucose, 25.0 NaHCO_3_, 2 sodium pyruvat, 3 myo inositol, 1 kynurenic acid. At room temperature slices equilibrated for at least 1 h in recording aCSF (in mM): 125 NaCl, 2.5 KCl, 1 MgSO_4_, 1.25 NaH_2_PO_4_, 2 CaCl_2_, 10 D-glucose, 25.0 NaHCO_3_, 2 sodium pyruvat, 3 myo inositol, 0.4 ascorbic acid, gassed with 95% O_2_ / 5% CO_2_, pH7.3.

#### Patch Clamp recordings

Coronal brain slices were placed in a submerged recording chamber mounted on an upright microscope (BX51WI, Olympus). Slices were continuously superfused with aCSF (2–3 ml/min, 32 °C, pH 7.3). Patch clamp recordings of miniature excitatory postsynaptic currents (mEPSCs) were performed as described previously (Sinning *et al.*, 2011). mEPSCs were recorded at a holding potential of −70 mV for at least 5 min in aCSF. Data analysis was performed off-line with the detection threshold levels set to 3 pA for mEPSCs. mEPSCs were isolated by adding tetrodotoxin (0.5 μM, Tocris Bioscience) and bicuculline methiodide (20 μM, Biomol) to block action potential-induced glutamate release and GABA_A_ receptor-mediated mIPSCs, respectively. 30 μM (2*R*)-amino-5-phosphonovaleric acid (dl-APV; Sigma-Aldrich) was added to suppress NMDA currents. The pipette solution contained the following (in mM): 120 CsMeSO_4_, 17.5 CsCl, 10 HEPES, 5 BAPTA, 2 Mg-ATP, 0.5 Na-GTP, 10 QX-314 [*N*-(2,6-dimethylphenylcarbamoylmethyl) triethylammonium bromide], pH 7.3, adjusted with CsOH. The following parameters were determined: frequency and peak amplitude.

### CPP, Diazepam and saline injections

To investigate the role of the N-methyl-D-aspartate (NMDA)-receptor on V1 responsiveness and OKR thresholds we administrated the competitive NMDA-receptor blocker (WD+CPP: n=8) (R, S)-3-(2-carbooxypiperazin-4-yl)propyl-1-phosphonic (CPP, Abcam). CPP was diluted in saline and injected intraperitoneally (i.p.) every 24 h at a dose of 12-15 mg/kg in a volume of 0.12 ml (Sato & Stryker, 2008; Teichert *et al.*, 2018b). To increase the level of cortical inhibition in WD mice for imaging experiments we intraperitoneally injected 0.12 ml diazepam solution (in saline, 1mg/kg; n=4) daily. In control mice (n=4), we daily injected the same volume of saline (i.p.).

### High performance liquid chromatography (HPLC)

Brain micropunches were taken from 1 mm V1 slices at −3.28 from Bregma from control (n=5) and WD mice (n=6) and homogenized by ultrasonication in 20 vol of 0.1 N perchloric acid at 4 °C immediately after collection. A total of 100 ml of the homogenate was added to equal volumes of 1 N sodium hydroxide for measurement of protein content. The remaining homogenate was centrifuged at 17 000 g and 4 °C for 10 min. Glutamate and GABA levels were determined using methods described previously (Winter *et al.*, 2009). Briefly, amino acids were precolumn-derivatized with o-phthalaldehyde-2-mercaptoethanol using a refrigerated autoinjector and then separated on a HPLC column (ProntoSil C18 ace-EPS) at a flow rate of 0.6 ml/min and a column temperature of 40 °C. The mobile phase was 50 mM sodium acetate (pH 5.7) in a linear gradient from 5% to 21% acetonitrile. Derivatized amino acids were detected by their fluorescence at 450 nm after excitation at 330 nm.

### Optomotor system

To determine subcortically mediated vision thresholds for spatial frequency and contrast of the optomotor response after WD (n=11), AD (n=4) or in untreated control mice (n=4), we used a virtual optomotor system (Prusky *et al.*, 2004). Briefly, placed on a platform, freely moving animals were surrounded by moving vertical sine wave gratings of varying spatial frequencies and contrasts. Mice reflexively track grating by head movements (optokinetic reflex, OKR) as long as they can see it (Prusky *et al.*, 2004). Thresholds for spatial frequencies were measured at 100% contrast and the contrast thresholds were determined at a spatial frequency of 0.2 cycles per degree (cyc/deg). From contrast thresholds contrast sensitivity was calculated (contrast sensitivity=(1/ contrast thresholds)×100)). We measured both spatial frequency and contrast sensitivity for each eye separately and, because they were almost identical, averaged these measurements for data presentation.

### V1 aspiration

To investigate whether V1 is required for the observed effects of WD on the OKR, the monocular and binocular V1 was bilaterally aspirated in WD (n=3) and control mice (n=3). First, mice received an injection of carprofene (5 mg/kg) for pain prevention. The correct position of V1 was determined using intrinsic imaging as described previously (Teichert & Bolz, 2017b). Briefly, to localize V1, animals were stimulated with a moving 2° wide horizontal light bar presented on the monitor placed in the right or left visual field at a distance of 25 cm to stimulate right and left eye, respectively. The bar covered 79° azimuth. The obtained retinotopic color coded phase map was then merged with a picture of the skull vascular pattern. Through a burr hole V1 was aspirated bilaterally as described previously (Prusky *et al.*, 2006) and the skin was re-sutured. Animals received subcutaneous carprofene (5 mg/kg) daily for pain prevention.

### Nissl staining

To demonstrate the efficiency of V1 aspiration experiments, we sacrificed mice tested in the Optomotry and performed a Nissl staining in the obtained brain slices. For this, brain sections were fixed in ethanol (95%) containing 5% acetic acid (99.5%) for 30 min. After washing with distilled water sections were incubated in a cresyl violet solution (0.1% in distilled water) for 4 min. After a further incubation in ethanol with ascending concentrations (50%, 70%, 95%, 99.9%) and xylol (98%), sections were embedded in depex (Serva). The sections were observed using a bright field microscope (Olympus) using a 10x objective.

### Experimental design and statistical analysis

To investigate whether WD or AD affect the responsiveness of the visual cortex we performed before-after comparisons of optical imaging data by paired *t*-tests and between-group comparisons by unpaired *t*-tests. Electrophysiological measurements and HPLC data as well were compared by an unpaired *t-test*. To examine potential effects of the cross-sensory deprivation on the visually mediated OKR, behavioral data of control, WD and AD mice (spatial frequency and contrast thresholds) were first analyzed by a one-way ANOVA with repeated measurements. After this group data were compared by *post hoc* two-tailed unpaired student’s *t-*test. The resulting *p*-values were then Bonferroni corrected. In the graphs, the levels of significance were set as: *p<0.05, **p<0.01, ***p<0.001. Data were analyzed using GRAPHPAD PRISM and SPSS and are presented as means and standard error of the mean (s.e.m) or as measurements of individual animals.

## Data availability statement

The dataset generated and analyzed within the present study are available from the corresponding author on reasonable request.

## Results

### Cross-modally induced shifts of ocular dominance

We here investigated the cross-modal effects of the deprivation of a non-visual sensory modality on visually evoked activity in the spared binocular V1 in fully adult mice adult mice (PD 120-240). For this, we induced either a somatosensory deprivation by bilaterally removing the macro-vibrissae (whisker deprivation, WD; n=7) (Teichert *et al.*, 2018a; Teichert *et al.*, 2018b) or an auditory deprivation (AD, n=6) by bilateral malleus removal (Teichert & Bolz, 2017b; Teichert *et al.*, 2017) and performed repeated intrinsic signal imaging experiments directly after either WD or AD (day 0) and 3 and 7 days after WD or AD (**Figure 1a**). Intrinsic signal imaging allows repeated non-invasive measurements of V1 responses evoked by visual stimulation (Kaneko *et al.*, 2008; Fu *et al.*, 2015) and its reliability has been profoundly validated by electrophysiological recordings (Kaneko *et al.*, 2008; Kaneko & Stryker, 2014). Since the binocular V1 of mice receives input of both eyes, we measured V1 activity evoked by visual stimulation of the contralateral and ipsilateral eye separately (**Figure 1b**). As a visual stimulus we used a drifting light bar of 100% contrast which was presented in the right binocular visual field (**Figure 1b**).

**Figure 1:**
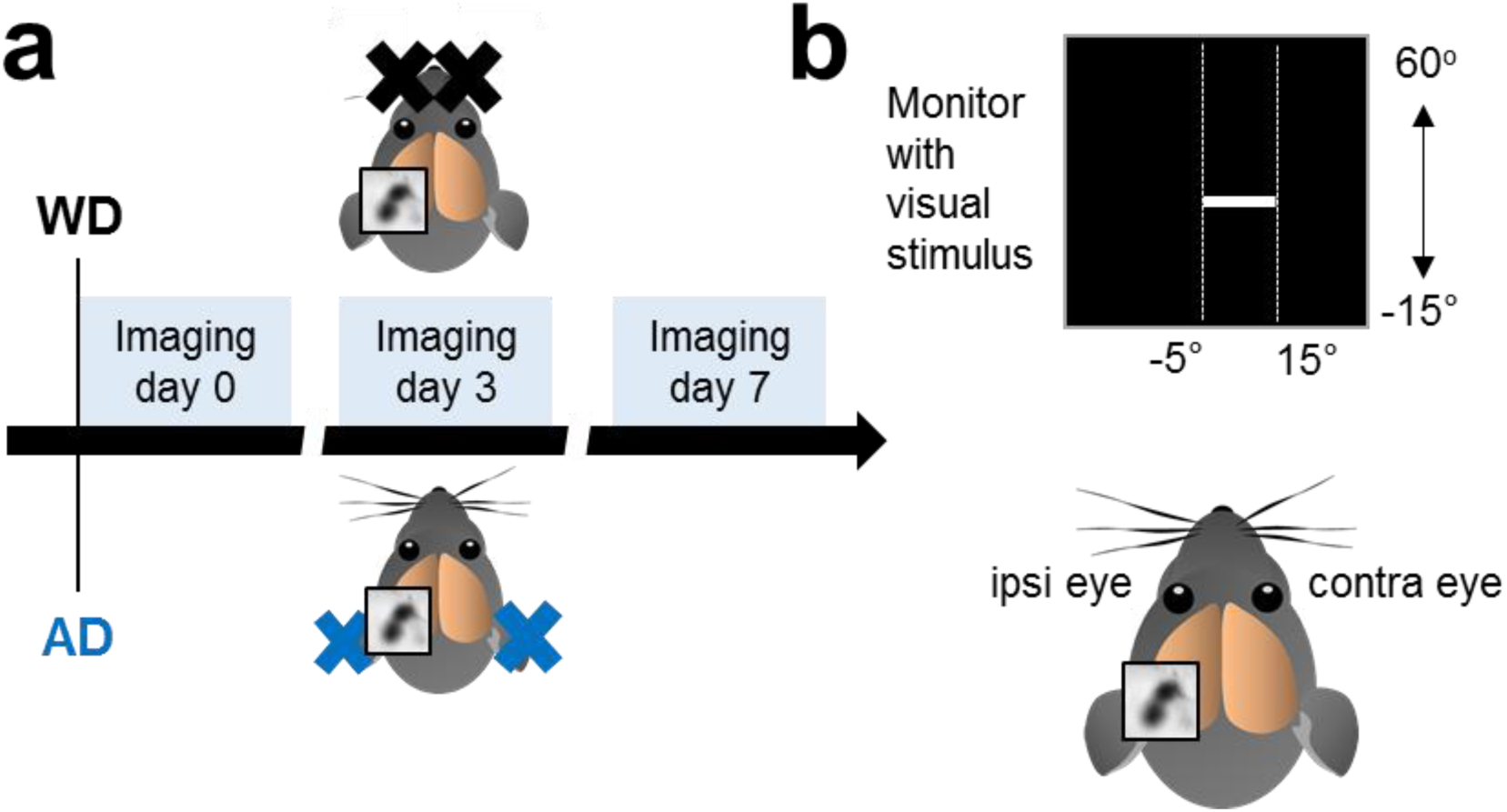
Experimental time course and schematic illustration of intrinsic signal imaging. (**a**) In one group of mice we performed WD by bilaterally plucking all macro-vibrissae and in another group we induced AD by bilateral malleus removal. The first imaging session (day 0) for mapping V1 started immediately after the deprivation followed by a second imaging session at day 3 and a third imaging session at day 7. (**b**) For visual stimulation we used an upward or downward moving white light bar with 100% contrast which was presented in the right binocular visual field. We always recorded V1 responses in the left hemisphere. Thus, according to the position of the recorded hemisphere the left eye represents the ipsilateral and the right eye represents the contralateral eye.

First, we examined whether repeated optical intrinsic imaging *per se* affects visually evoked V1 activity in normal untreated mice (n=5). **Figure 2a** depicts representative V1 activity maps evoked by the stimulation of either the contralateral (upper row) or the ipsilateral eye (lower row) obtained at 0, 3 and 7 days. Darker activity maps indicate higher visually driven V1 responses. It is clearly visible that V1 input strength remained unchanged during the time tested, with the contralateral eye always dominating the input to V1, the normal OD ratio for the binocular region of V1 (Drager, 1975). These results demonstrate that repeated intrinsic imaging provides reliable and stable measurements of sensory evoked V1 activity.

**Figure 2:**
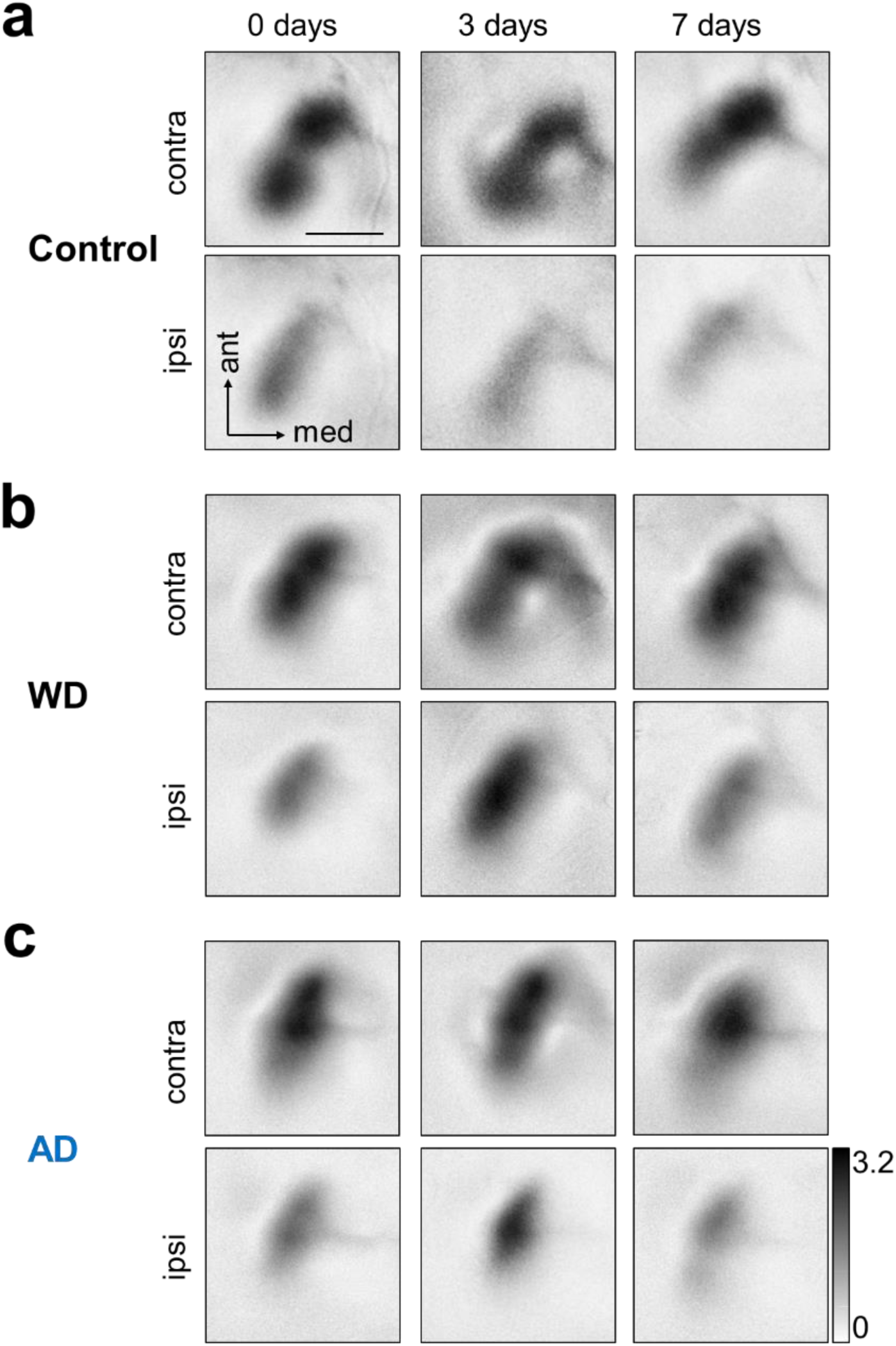
Representative V1 maps evoked by the stimulation of either the contralateral or ipsilateral eye at day 0, 3 and after 7 days. (**a**) Upper row: In normal control mice contralateral eye stimulation always evoked similarly strong V1 responses. Lower row: V1 responses evoked by ipsilateral eye stimulation were equally strong over the time tested but these responses were always weaker that V1 maps obtained after visual stimulation of the contralateral eye. (**b**) Upper row: Like in control mice, WD did not lead to alterations of V1 responses elicited by contralateral eye input. Lower row: However, V1 responses elicited by ipsilateral eye stimulation markedly increased 3 days after WD which was followed by a decrease of V1 response strength 7 days after WD. (**c**) Upper row: Responsiveness of V1 to visual stimulation of contralateral eye remained stable after AD over the time tested. Lower row: 3 days after AD there was a massive increase of V1 responses evoked by ipsilateral eye stimulation which was followed by a recovery of V1 activity 7 days after AD. Thus, the deprivation in both senses (somatosensory or auditory) alters the OD within the binocular zone of V1. Scale bar: 1 mm

Next, we tested whether WD cross-modally affects responsiveness of V1 to contra- or ipsilateral eye stimulation. As shown in **Figure 2b** (upper row) V1 activity patches elicited by visual stimulation of the contralateral eye remained equally strong at 0, 3 and 7 days after WD. However, surprisingly, there was a marked increase of V1 responses evoked by ipsilateral eye stimulation 3 days after WD which was followed by a decrease of V1 input strength back to the level of the V1 maps obtained at 0 days. These results suggest that WD provokes a transient shift of OD within the binocular zone of V1 after 3 days, which during the following 4 days then readjusted to the original level, probably due to homeostatic mechanisms.

We further investigated whether the deprivation of another non-visual modality, the auditory sense, induces similar cross-modal effects in V1. **Figure 2c** shows representative V1 maps evoked by the stimulation of either the contralateral (upper row) or ipsilateral eye (lower row) obtained at 0, 3 and 7 days after AD. V1 maps elicited by visual stimulation of the contralateral eye remained stable over the whole time tested. However, like already found after WD, V1 response maps driven by the ipsilateral eye were markedly stronger 3 days after AD. This increase of elicited V1 activity was followed by a decrease back to starting levels 7 days after AD. Thus, our results indicate that the deprivation of non-visual sensory modalities leads to cross-modal alterations of OD. Notably, this took place without monocular deprivation, the common traditional paradigm to induce OD shifts in mammals used up to now since its first description 55 years ago (Wiesel & Hubel, 1963; Hubel & Wiesel, 1970; Gordon & Stryker, 1996)

Quantification revealed that repeated intrinsic signal imaging of untreated control animals at 0, 3 and 7 days did not influence V1 responsiveness after contra-or ipsilateral eye stimulation (contra: 0 d vs 3 d: *p*=0.97; 0 d vs 7 d: *p=0.77*; 3 d vs 7 d: *p*=0.83; ipsi: 0 d vs 3 d: *p*=0.34; 0 d vs 7 d: *p=0.24*; 3 d vs 7 d: *p*=0.40; paired *t*-tests; **Figure 3a, b; Table 1**). Hence, the ocular dominance index (ODI) in these animals remained stable during the time tested (0 d vs 3 d: *p*=0.79; 0 d vs 7 d: *p=0.47*; 3 d vs 7 d: *p*=0.53; paired *t*-tests; **Figure 3c; Table 1**), indicating that this technique provides stable data of visually evoked V1 activity.

**Table 1:**
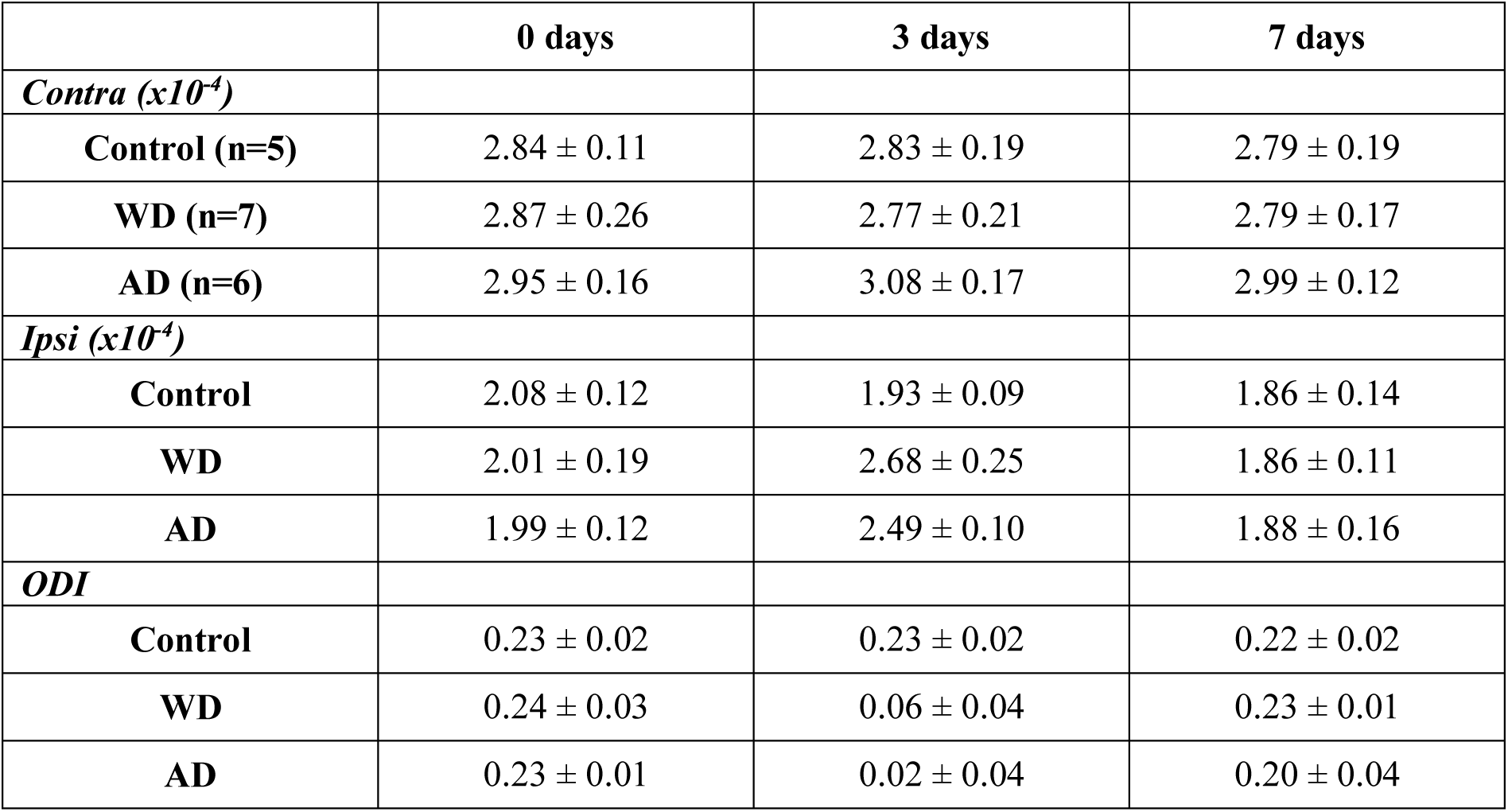
The effects of WD and AD on V1 responsiveness and OD. Data are presented as means ± s.e.m.

**Figure 3:**
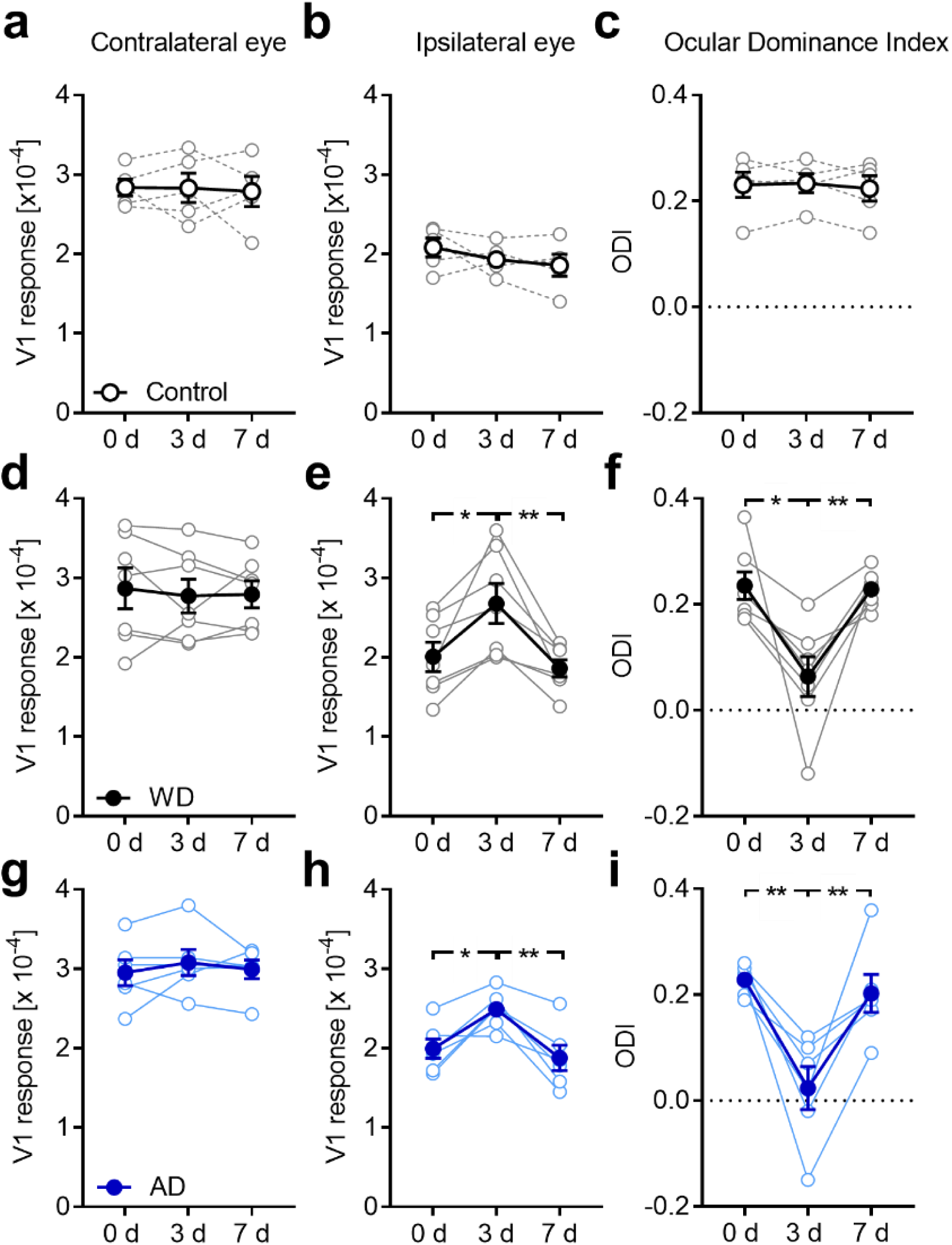
Both WD and AD shift OD in the binocular V1 in fully adult mice. (**a, b**) V1 activity evoked by visual stimulation of the contra or ipsilateral eye in untreated control mice (n=5) remained unchanged at 0, 3 and 7 days. (**c**) Thus, over the same time course the ODI did not change underlining the reliability of repeated intrinsic signal imaging. (**d**) During one week after WD (n=7), V1 activity elicited by contralateral eye stimulation was unaltered. (**e**) However, V1 responsiveness to ipsilateral eye stimulation markedly increased 3 days after WD followed by a recovery of V1 activity 7 days after WD. (**f**) These alterations of V1 responsiveness led to an ODI shift towards zero at day 3 which was followed by a readjustment of the ODI 7 days after WD. (**g**) After AD (n=6) V1 activity elicited by contralateral eye stimulation remained unchanged during the time tested. (**h**) However, V1 responses evoked by visual stimulation of the ipsilateral eye massively increased after 3 days of AD. After the 7 days V1 responses due to the ipsilateral eye input decreased back to the starting level measured at day 0. (**i**) Hence, the ocular dominance index displayed a dramatic shift towards zero 3 days after AD which was followed by a complete recovery after one week. Thus, the deprivation of a non-visual input altered OD in the spared V1. Open circles represent measurements of individual animals. Closed circles represent the means of each group ± s.e.m.; *p<0.05, **p<0.01

After WD, V1 activation elicited by contralateral eye stimulation did not change during the first week (0 d vs 3 d: *p* =0.54; 0 d vs 7 d: *p=*0.55; 3 d vs 7 d: *p* =0.87; paired *t*-test; **Figure 3d; Table 1**). However, cortical responses elicited by ipsilateral eye stimulation significantly increased 3 days after WD (0 d vs 3 d: p=0.01; paired *t-*test; **Figure 3e; Table 1**) followed by a significant decrease back to starting levels after another 4 days (3 d vs 7 d: *p=*0.007; 0 d vs 7 d, *p=*0.36; paired *t-*tests; **Figure 3e; Table 1**). Thus, over the same time course there was a remarkable drop of the ODI towards zero which was followed by a complete recovery at 7 days after WD (0 d vs 3 d: *p=*0.003; 3 d vs 7 d: *p=*0.005; 0 d vs 7 d: *p=*0.77; paired *t*-tests; **Figure 3f; Table 1**). These data indicate that WD leads to an increased V1 activation to visual stimulation after 3 days which is, however, restricted to the ipsilateral eye input. Again, the restoration of V1 activity and ODI after 7 days might suggest a homeostatic mechanism adjusting V1 inputs back to baseline levels.

Quantification of the cross-modal effects of AD on visually evoked V1 activity reveled that V1 responsiveness to contralateral eye stimulation was unchanged at 0, 3 and 7 days after AD (0 d vs 3 d: *p*=0.31; 0 d vs 7 d: *p=0.84*; 3 d vs 7 d: *p*=0.50; paired *t*-tests; **Figure 3g; Table 1**). However, V1 activity elicited by visual stimulation of the ipsilateral eye dramatically increased 3 days after AD (0 d vs 3 d: *p*=0.01; paired *t*-tests; **Figure 3h; Table 1**) and decreased again after 7 days of AD to the level obtained at 0 days (0 d vs 7 d: *p=0.42*; 3 d vs 7 d: *p*=0.005; paired *t*-tests; **Figure 3h; Table 1**). As a direct consequence of the observed V1 activity changes there was a dramatic shift of the ODI towards zero 3 days after AD, which was followed by a full recovery of OD (0 d vs 3 d: *p*=0.005; 0 d vs 7 d: *p=0.56*; 3 d vs 7 d: *p*=0.009; paired *t*-tests; **Figure 3i; Table 1**).

Group comparison showed that at day 0 V1 activity evoked by either the contra or ipsilateral eye and the ODI was almost identical in control, WD and AD animals (contra: control vs WD: *p=*0.94, control vs AD: *p=*0.60; ipsi: control vs WD: *p=*0.75, control vs AD: *p=*0.62; ODI: control vs WD: *p=*0.91, control vs AD: *p=*0.92; unpaired *t*-tests; **Figure 4a, b, c; Table 1**). These results suggest that V1 responsiveness to the separate stimulation of both eyes is unchanged immediately after WD or AD. Comparison of data obtained at day 3 showed that V1 responses elicited by visual stimulation of the contralateral eye were similar in control, WD and AD mice (control vs WD: *p*=0.84; control vs AD: *p*=0.37; unpaired *t*-tests; **Figure 4d; Table 1**). However, V1 activity evoked by ipsilateral eye stimulation was significantly increased after both WD and AD compared to controls (control vs WD: *p*=0.03; control vs AD: *p*=0.002; unpaired *t*-tests; **Figure 4e; Table 1**). Hence, ODI values were significantly smaller in WD and AD mice (control vs WD: *p*=0.005; control vs AD: *p*=0.002; unpaired *t*-tests; **Figure 4f; Table 1**). Group comparison at 7 days revealed that V1 responses elicited by visual stimulation of the contralateral in both WD and AD mice were similar to control animals (control vs WD: *p*=0.99; control vs AD: *p*=0.37; unpaired *t*-tests; **Figure 4g; Table 1**). Moreover, after 7 days evoked V1 amplitudes due to ipsilateral eye stimulation as well as the ODI in WD or AD animals were completely restored back to control levels (ipsi: control vs WD: *p*=0.99; control vs AD: *p*=0.93; ODI: control vs WD: *p*=0.86; control vs AD: *p*=0.64; paired t-tests; **Figure 4h, i; Table 1**).

**Figure 4:**
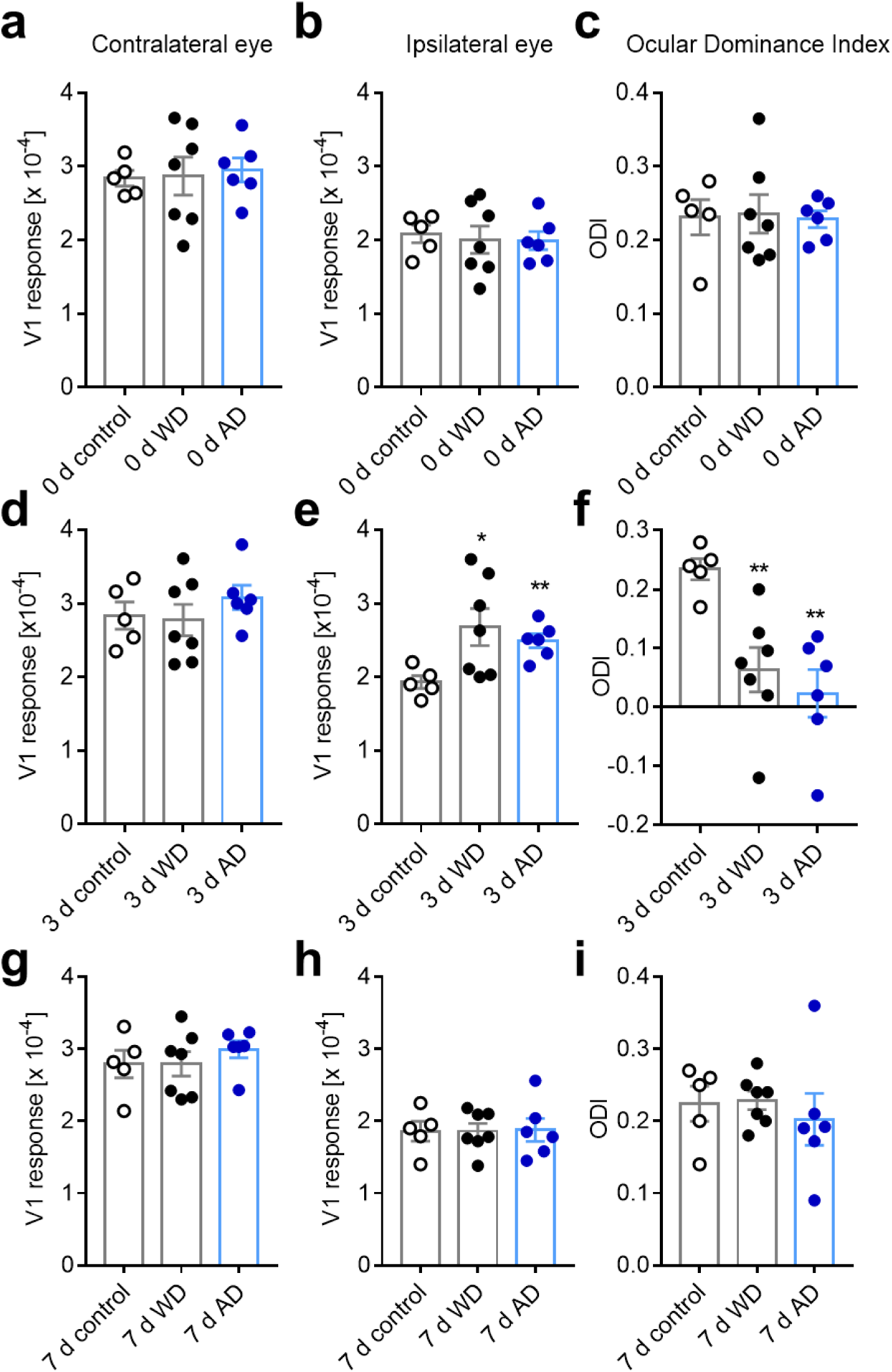
Both WD or AD induce an ODI shift compared to normal control mice. (**a, b**) V1 responses evoked by visual stimulation of either the contra or ipsilateral eye immediately after WD (n=7) or AD (n=6) were indistinguishable from values obtained in normal control mice (n=5). (**c**) Hence, directly after WD or AD there was no alteration of the ODI. (**d**) After 3 days of either WD or AD V1 responses elicited by contralateral eye stimulation was not different from control values. (**d**) However, V1 activity evoked by stimulation of the eye ipsilateral to the recorded hemisphere was dramatically increased 3 days after WD or AD compared to control levels. (**f**) Thus, at this time point there was a highly significant shift of the ODI towards zero. (**g**) After 7 days of WD or AD V1 responses elicited by the stimulation of contralateral eye were unchanged compared to the values of control mice. (**h**) Interestingly, V1 responsiveness to ipsilateral eye stimulation was re-adjusted to control levels after one week of either WD or AD. (**i**) Consequently, the ODI of WD or AD mice was completely restored back to control values after 7 days. Bars represent the means ± s.e.m. and open or filled circles represent measurements of individual animals; *p<0.05, **p<0,01

Taken together, our data indicate that the deprivation of a non-visual sense leads to a selective increase of V1 activity evoked by the typically “weaker”, ipsilateral eye and thus, to changes of the OD within the binocular zone of V1. Thus, these results suggest that sensory deprivations of non-visual sensory modalities can induce neuronal plasticity in spared primary sensory cortices.

### Cross modally induced changes of OD cannot be explained by a saturation of V1 activity evoked by the contralateral eye

So far we describe that both WD and AD lead to a marked ODI shift which was mediated by a selective increase of V1 activity evoked by ipsilateral eye stimulation after 3 days, whereas the contralateral eye input to V1 remains unchanged. Since we were surprised by this unexpected selective cross-modal effect, we wondered whether the absence of V1 activity changes due to contralateral eye stimulation 3 days after deprivation might be caused by a saturation of the contralateral eye input to V1. To address this issue we first investigated whether V1 activity elicited by contralateral eye stimulation is already saturated in normal control mice (n=4). For this, we measured V1 responses evoked by monocular ipsilateral and contralateral eye stimulation and also after binocular visual stimulation (**Figure 5a**). As expected, we found that V1 responses evoked by ipsilateral eye stimulation responses were always weaker than after stimulation of the contralateral eye (ipsi vs contra: 2.31±0.15 (×10^−4^) vs 3.18±0.11 (×10^−4^), *p=*0.008; paired *t-*test; **Figure 5b**). However, V1 activity evoked by contralateral eye stimulation was significantly weaker than after binocular stimulation (contra vs bino: 3.18±0.11 (×10^−4^) vs 3.87±0.11 (×10^−4^), *p*=0.003; paired *t*-tests; **Figure 5b**). Thus, these data indicate that V1 responsiveness, as measured by intrinsic signal imaging, is not saturated after contralateral eye stimulation.

**Figure 5:**
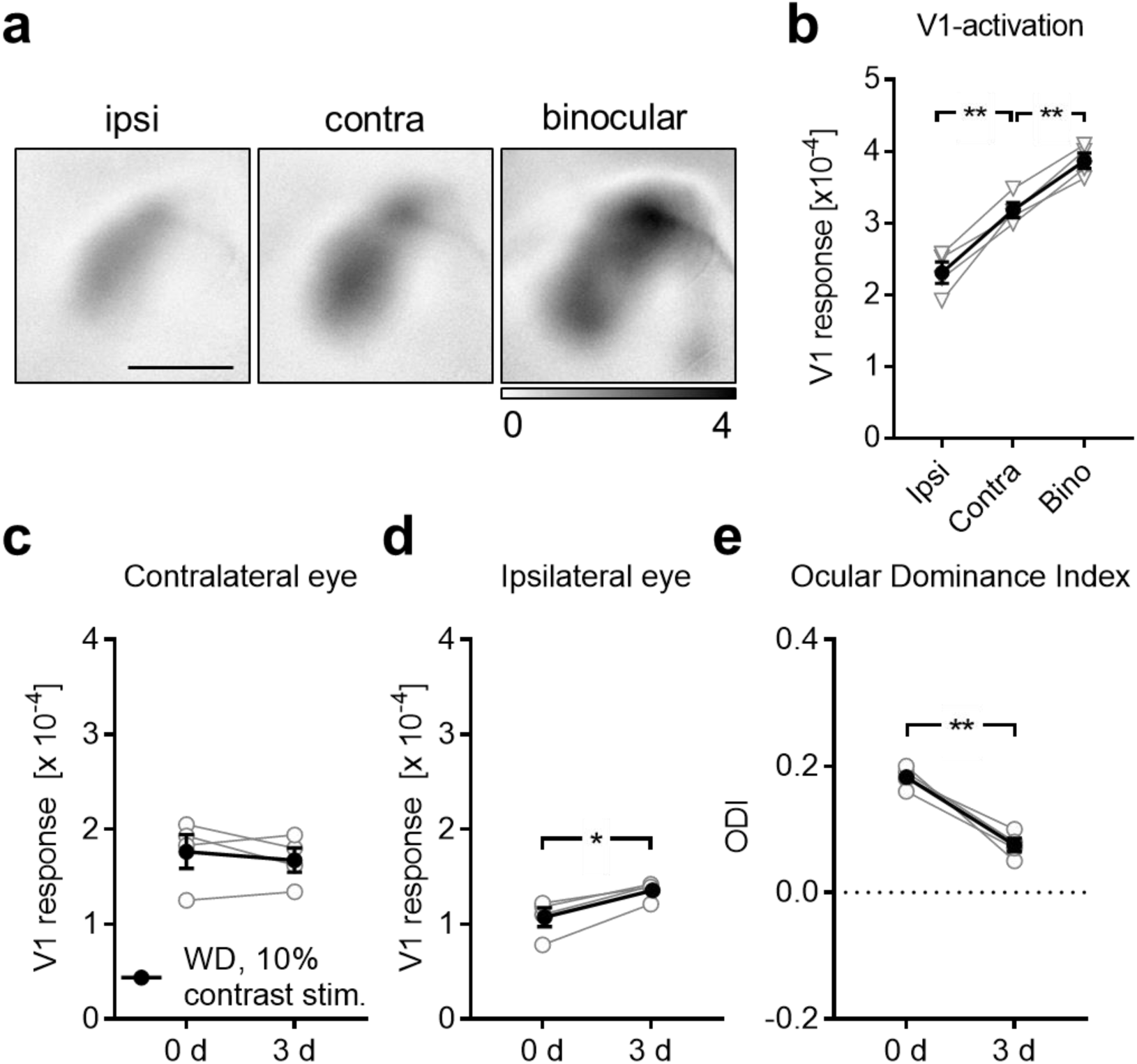
Exclusion of saturation of V1 responses evoked by contralateral eye stimulation: (**a**) Representative V1 amplitude maps evoked by visual stimulation of the ipsi and contralateral eye and elicited by binocular stimulation of normal untreated mice (n=4). (**b**) V1 responsiveness to ipsilateral eye stimulation was always weaker compared to V1 responsiveness to contralateral eye stimulation. However, V1 activity elicited by visual stimulation of the contralateral eye was significantly weaker than V1 activity evoked by binocular stimulation. Hence, V1 responses, measured with intrinsic signal imaging, are not saturated by the input through the contralateral eye. (**c**) V1 responses evoked by contralateral eye stimulation with a 5% contrast visual stimulus at 0 and 3 days after WD (n=4) remained unchanged. (**d**) However, there was a potentiation of V1 responses to the input through the ipsilateral eye between 0 and 3 days after WD. (e) Thus, the ODI significantly shifted towards zero. Hence, visual stimulation with a weaker visual stimulus reveals the same effect of WD on V1 activity like visual stimulation with a strong visual stimulus. Open circles represent measurements of individual animals. Closed circles represent the means of each group ± s.e.m.; *p<0.05, **p<0.01; Scale bar: 1 mm

To further exclude a potential saturation effect, we again measured V1 responsiveness to contra and ipsilateral eye stimulation 0 and 3 days after WD (n=4). However, this time we reduced the contrast of the visual stimulus from 100% to 10%, which generally decreases visually evoked V1 responses (Teichert & Bolz, 2017b; Teichert *et al.*, 2018a; Teichert *et al.*, 2018c). Hence, potential changes of V1 input from the contralateral eye 3 days after WD can be detected. Quantification showed that V1 activity elicited by visual stimulation of the contralateral eye still remained unchanged between 0 and 3 days after WD (0 d vs 3 d: 1.77±0.18 (×10^−4^) vs 1.68±0.13 (×10^−4^), *p*=0.47; paired *t-*test; **Figure 5c**) whereas ipsilateral eye input to V1 significantly increased again (0 d vs 3 d: 1.07±0.10 (×10^−4^) vs 1.36±0.05 (×10^−4^), *p*=0.01; paired *t-*test; **Figure 5d**). Thus, the differential V1 activity changes caused a marked reduction of the ODI 3 days after WD (0 d vs 3 d: 0.18±0.009 vs 0.08±0.01, *p=*0.006; paired *t-*test; **Figure 5e**). These data suggest that cross-modally induced OD changes in V1 are independent of the strength of the presented visual stimulus. Hence, our results make it reasonable to conclude that the absence of V1 activity changes evoked by contralateral eye stimulation 3 days after WD is not caused by a saturation of the contralateral eye input to V1.

### Cross-modally induced changes of ocular dominance require visual experience

It has been suggested that plastic changes in a spared primary sensory cortex require sensory experience through its main input (Petrus *et al.*, 2014). Thus, we next asked whether patterned visual input through the contra or ipsilateral eye is necessary to provoke V1 activity changes 3 after WD. To address this question we first combined WD with MD of the contralateral eye (n=4) and measured V1 responsiveness at 0 and 3 days using intrinsic signal imaging. We found that V1 responses evoked by visual stimulation of the contralateral (closed) eye remained unchanged whereas V1 activity driven by the ipsilateral (open) eye input was increased 3 days after WD and MD (contra: 0 d vs 3 d: 2.79±0.13 (×10^−4^) vs 2.81±0.14 (×10^−4^), *p=*0.86; ipsi: 0 d vs 3 d: 1.83±0.06 (×10^−4^) vs 2.30±0.08 (×10^−4^), *p*=0.002; paired *t-*tests; **Figure 6a, b**). These activity alterations led again to a significant reduction of the ODI (0 d vs 3 d: 0.24±0.03 vs 0.09±0.03, *p=*0.009; paired *t-*test; **Figure 6c**), similar to WD mice with open contralateral eyes. Thus, cross-modally induced changes of OD do not require patterned visual input through the contralateral eye.

**Figure 6:**
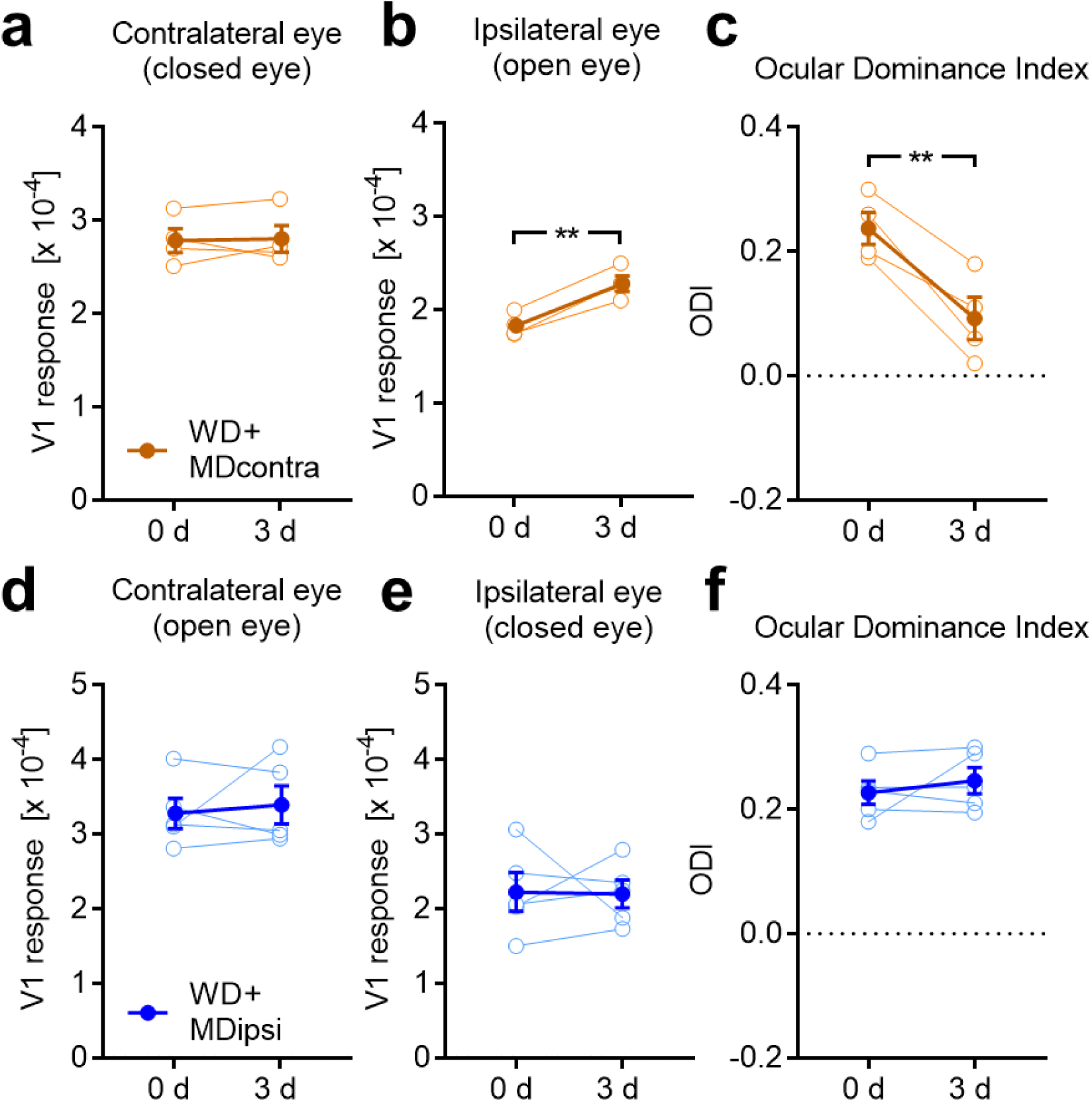
Cross-modally induced ODI shift requires patterned vision through the ipsilateral eye. (**a**) Combining WD with a MD of the contralateral eye (n=4) did not lead to changes of V1 responses evoked by the contralateral eye between 0 and 3 days after WD. (**b**) However, there was a significant increase of V1 activity elicited by visual stimulation of the ipsilateral eye, like found after WD only. (**c**) Thus, the ODI markedly shifted towards zero. (**d, e**) In contrast, if we combined WD with a MD of the ipsilateral eye (n=5), V1 activity evoked by both contra and ipsilateral eye stimulation remained statistically unchanged after 3 days. (**f**) Moreover, there was no ODI shift after this treatment suggesting that patterned vision through the ipsilateral eye is required for cross-modally induced OD shifts. Open circles represent measurements of individual animals. Closed circles represent the means of each group ± s.e.m.; *p<0.05, **p<0.01

Next, we investigated whether experience of patterned vision through the ipsilateral eye is required for WD induced changes of OD. Combining WD with a MD of the ipsilateral eye (n=5) did not lead to changes of the contralateral eye input to V1 (0 d vs 3 d: 3.28±0.20 (×10^−4^) vs 3.40±0.25 (×10^−4^), *p=*0.67; paired *t*-test; **Figure 6d**). However, this treatment abolished the selective increase of V1 responsiveness to ipsilateral eye stimulation after 3 days of WD (0 d vs 3 d: 2.23±0.26 (×10^−4^) vs 2.20±0.19 (×10^−4^), *p=*0.94; paired *t-*test; **Figure 6e**). Hence, the ODI did not change after these interventions (0 d vs 3 d: 0.23±0.02 vs 0.25±0.02, *p=*0.45; paired *t-*test; **Figure 6f**). These results suggest that cross-modal changes of V1 activity, observed after WD, depend on visual experience selectively through the eye ipsilateral to the recorded hemisphere.

### WD cross-modally increases mEPSC amplitudes in V1 layer 4

It has been demonstrated that dark exposure cross-modally potentiates layer 4 synapses in the spared A1 (Petrus *et al.*, 2014). Hence, we here examined the cross-modal effects of WD on the strength of layer 4 synapses in the spared V1. For this, we performed whole-cell recordings in acute V1 slices of normal control mice (11 cells, n=4 mice) and animals 3 d after WD (6 cells, n=3 mice). We did not find changes in frequency of miniature excitatory postsynaptic currents (mEPSC) after WD (control: 2.42±0.85 (Hz), 3 d WD: 3.19±0.93 (Hz), *p*=0.49; unpaired *t-*test). However, there was a significant increase in α-amino-3-hydroxy-5-methyl-4-isoxazolepropionic acid receptor (AMPAR) mediated mEPSC amplitudes (control: 9,31±0.58 (pA), 3 d WD: 15.02±1.63 (pA), *p=*0.0011; unpaired *t*-test; **Figure 7a, b**) indicating that WD, indeed, cross-modally increased the strength of excitatory V1 layer 4 synapses, which may include thalamo-cortical synapses driven by the ipsilateral eye. Strengthening of thalamo-cortical synapses typically leads to an increased sensory driven responsiveness of primary sensory cortices (Heynen & Bear, 2001; Petrus *et al.*, 2014). Consistent with these observations is our finding that V1 responses evoked by ipsilateral eye stimulation were increased 3 d after WD, as revealed by intrinsic imaging. Taken together, these data suggest that WD cross-modally induces synaptic plasticity in the spared V1.

**Figure 7:**
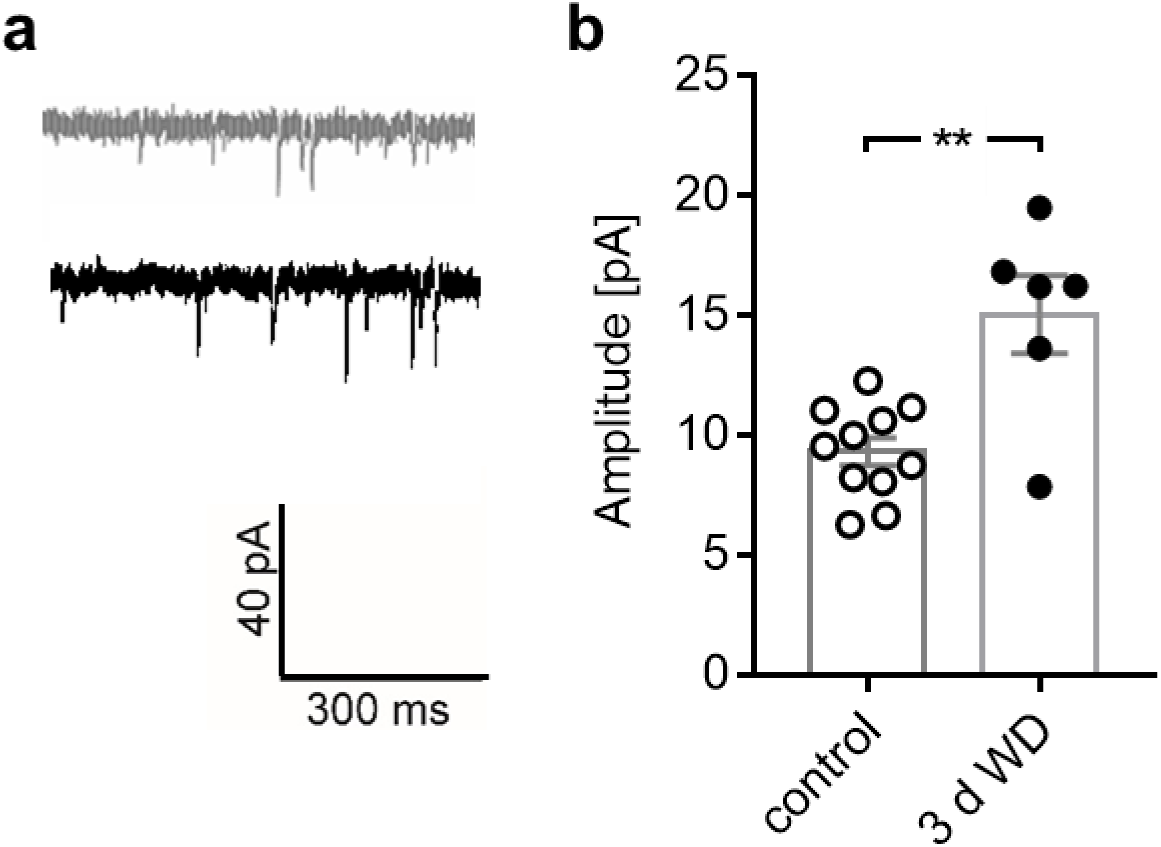
WD cross-modally increases mEPSC amplitudes in V1 layer 4. (**a**) Representative traces of mEPSCs recorded in control mice (n=4) and 3 d after WD (n=3). (b) The amplitude of mEPSCs was significantly increased in WD mice. These results suggest that WD induces synaptic plasticity in V1 layer 4. Bars represent means together with s.e.m., Open and filled circles represent measurements of individual animals; **p<0.01

### WD cross-modally increases the E/I ratio in V1

Experience dependent V1 plasticity that after MD leads to changes in OD typically declines with aging and is completely absent in fully adult mice (Lehmann & Lowel, 2008), like used in the present study. However, previous studies could demonstrate that increasing the cortical excitation/inhibition (E/I) ratio is a central hub for the restoration of visual plasticity in the adult V1 (He *et al.*, 2006; Sale *et al.*, 2007; Maya-Vetencourt *et al.*, 2008; Harauzov *et al.*, 2010). Hence, as a next step, we examined whether WD for 3 days leads to cross-modal changes in V1 glutamate and GABA levels. For this, we quantified levels of glutamate and GABA by post-mortem HPLC analyzes of V1 tissue from control mice (n=5) and WD mice 3 days after WD (n=6). Quantification showed that there was a significant increase in V1 glutamate levels 3 days after WD (control vs 3 d WD: 71.07±1.5 (nMol/mg protein) vs 77.70±1.79 (nMol/mg protein), *p=*0.02; unpaired *t*-test; **Figure 8a**). Moreover, V1 GABA content slightly decreased by about 6% after WD, which was, however, not statistically significant (control vs 3 d WD: 9.80±0.19 (nMol/mg protein) vs 9.26±0.39 (nMol/mg protein), *p*=0.27; unpaired *t*-test; **Figure 8b**). Due to the differential regulations of glutamate and GABA levels in V1 after WD, the glutamate/GABA ratio significantly increased 3 days after WD (control vs 3 d WD: 7.25±0.12 vs 8.11±0.30, *p=*0.02; **Figure 8c**). These results suggest that WD cross-modally alters the V1 E/I balance in favor of excitation, which might set the adult V1 back into a plastic stage.

**Figure 8:**
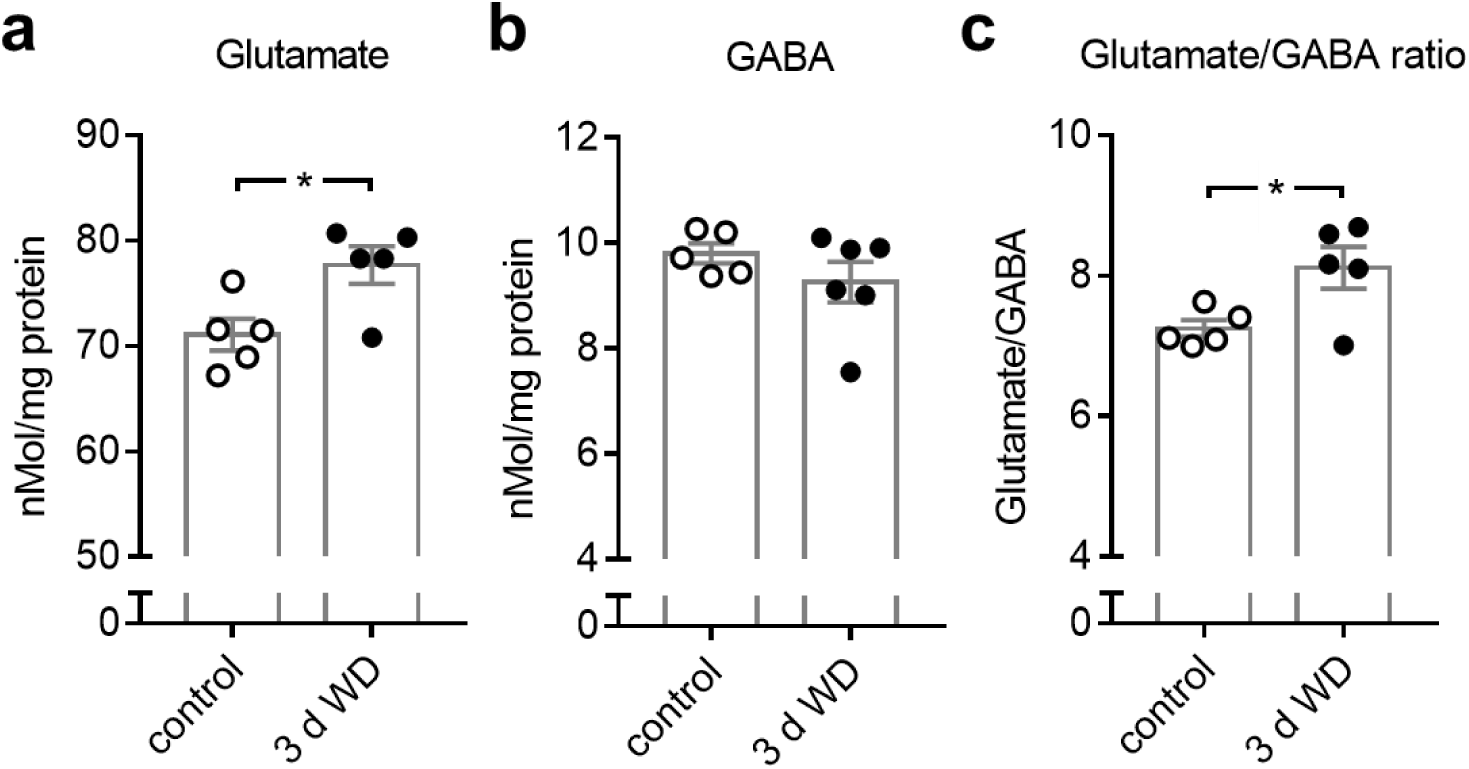
Concentration changes of neurotransmitters in V1 after WD revealed by post-mortem HPLC analyzes. (**a**) Compared to the V1 glutamate level of control mice (n=5), there was a significant increase of V1 glutamate content 3 days after WD (n=5). (**b**) There was slight but not significant reduction of the V1 GABA concentration at 3 days after WD (n=6) compared to controls. (**c**) The glutamate/GABA ratio was markedly increased at 3 after WD. Bars represent means together with s.e.m., Open and filled circles represent measurements of individual animals; *p<0.05

### Cross-modal changes of V1 activity depend on V1 GABA levels and NMDA receptor activation

We next investigated whether the observed increase of the V1 glutamate/GABA ratio after WD was related to cross-modally induced V1 activity changes. For this, we artificially raised cortical GABAergic inhibition by daily systemic administration of diazepam in WD mice (n=4) and measured V1 responsiveness at 0 and 3 days using intrinsic imaging. Diazepam, administrated systemically or locally, is a common tool for enhancing cortical inhibition, since it increases GABA receptor mediated currents (Spolidoro *et al.*, 2011; Greifzu *et al.*, 2014; Stodieck *et al.*, 2014). In control animals (n=4) we also performed WD and systemically administrated saline. As expected, saline treatment did not influence the cross-modal effects of WD on V1, as V1 activity evoked by visual stimulation of the contralateral eye was unchanged during the time tested, whereas the ipsilateral eye input significantly increased at day 3 after WD (contra: 0 d vs 3 d: *p=*0.38; ipsi: 0 d vs 3 d: *p=*0.006; paired *t-*tests; **Figure 9a, b; Table 2**). Consequently, ODI significantly decreased after WD (0 d vs 3 d: *p=*0.002; paired *t-*tests; **Figure 9c; Table 2**). However, in WD animals treated with diazepam both the contralateral and ipsilateral eye input strength to V1 and thereby also the ODI remained unchanged at 3 days (contra: 0 d vs 3 d: *p=*0.36; ipsi: 0 d vs 3 d: *p=*0.61; ODI: 0 d vs 3 d: *p=*0.89; paired *t-*tests; **Figure 9a, b, c; Table 2**). Thus, increasing cortical inhibition abolished the WD induced activity changes in V1. These results indicate that WD induced increase of the V1 E/I ratio plays an important role for cross-modally induced V1 plasticity.

**Table 2:**
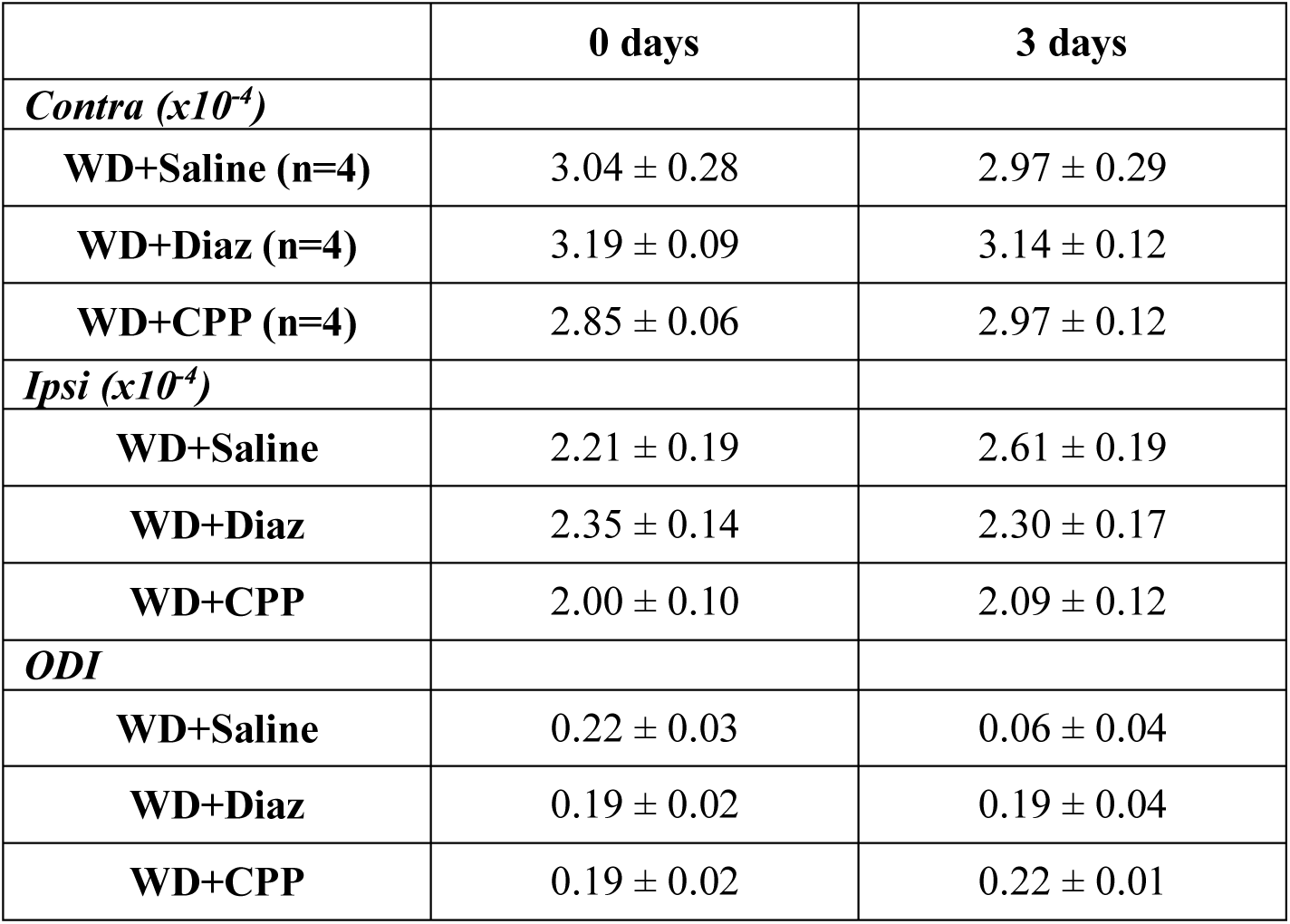
The effects of diazepam and CPP administration on cross-modally induced V1 activity changes after WD. Data are presented as means ± s.e.m.

**Figure 9:**
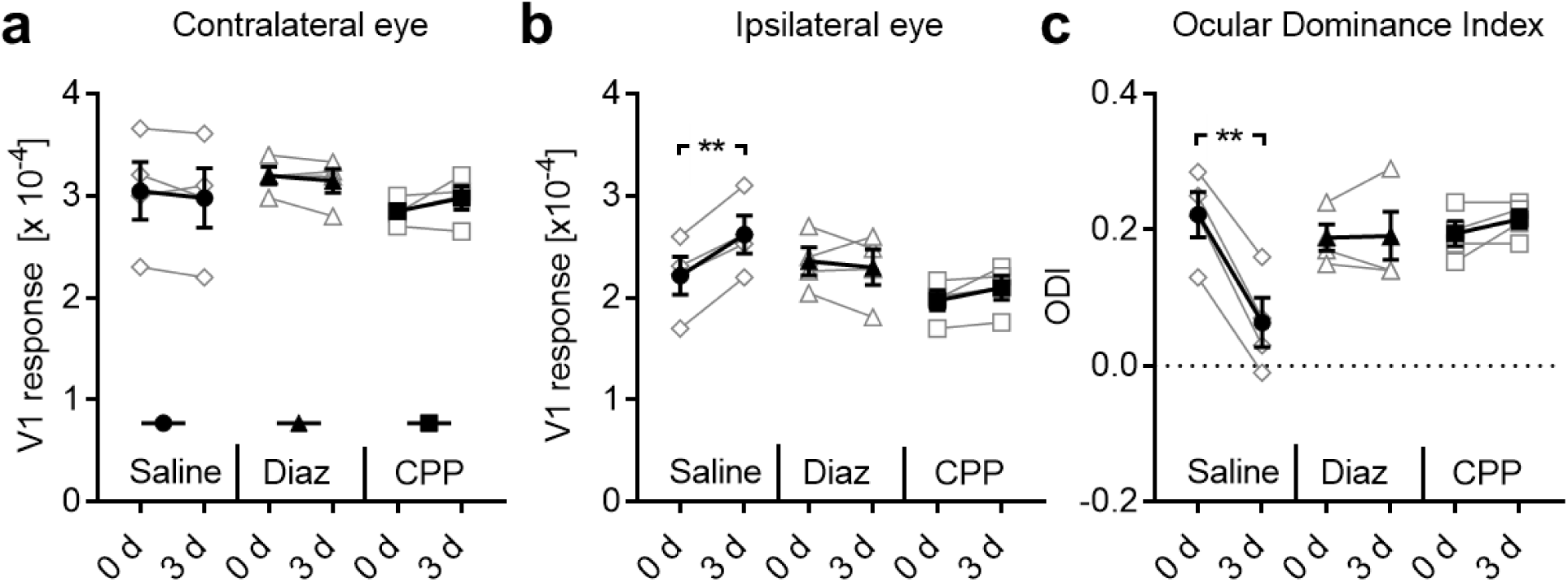
Both increasing inhibition and blocking NMDA receptor activation block cross-modally induced V1 plasticity. (**a**) In WD mice treated with saline (n=4) or diazepam (n=4) or CPP (n=4) V1 responses evoked by visual stimulation of the contralateral eye remained unchanged. (**b**) There was a significant increase of V1 responses elicited by ipsilateral eye stimulation in WD+Saline mice. However, these changes were completely abolished by diazepam or CPP injections. (**c**) We found a significant reduction of ODI in saline treated WD mice, whereas ODI did not change after diazepam of CPP administration. Taken together, our data suggest that cross-modally induced alterations of V1 OD depend on increased glutamateric excitation and NMDA receptor activation. Open circles represent measurements of individual animals. Closed circles represent the means of each group ± s.e.m.; **p<0.01

Next, we tested the hypothesis that WD induced V1 activity changes might rely on NMDA receptor (NMDAR) activation. Previous investigations could demonstrate an involvement of NMDARs in experience dependent V1 plasticity, as systemic administration of the competitive NMDA receptor antagonist CPP or genetic deletion of cortical NMDARs abolished plastic alterations in V1 (Sawtell *et al.*, 2003; Sato & Stryker, 2008). Moreover, we could recently show that blocking NMDAR activation by systemic administration of CPP abolished cross-modally induced restoration of ocular dominance plasticity (Teichert *et al.*, 2018b). Hence, WD mice received daily injections of CPP (n=4) and we measured V1 responsiveness again at 0 and 3 days. V1 responsiveness to contralateral eye stimulation as well as V1 activity elicited by visual stimulation of the ipsilateral eye did not change during the time tested (contra: 0 d vs 3 d: *p=*0.25; ipsi: 0 d vs 3 d: *p=*0.14; paired *t-*tests; **Figure 9a, b; Table 2**). Thus, the ODI remained unchanged in these mice (0 d vs 3 d: *p=*0.21; paired *t-*tests; **Figure 9c, Table 2**). These data show that systemic administration of CPP blocks WD induced cross-modal plasticity in V1. Taken together, our results suggest that NMDA receptor activation is necessary to provoke cross-modal strengthening of sensory driven activity in spared sensory cortices.

### WD cross-modally improves visual performance

As a next step we investigated whether cross-modal V1 response alterations are also reflected at the level of visually mediated behavior. In a recent study we could already demonstrate that WD markedly refined V1 mediated visual performance as revealed by visual water task experiments (Teichert *et al.*, 2018a). Another example of visual behaviors is the so called optokinetic reflex (OKR), a head and eye movement, mediated by subcortical structures, which stabilizes images on the retina (Liu *et al.*, 2016). Interestingly, previous studies could show that V1 activity can modulate the OKR (Prusky *et al.*, 2006; Liu *et al.*, 2016). We therefore hypothesized that the observed cross-modally induced changes of visually driven V1 activity (after WD or AD) might also lead to changes of the OKR. Hence, we next investigated the repercussions of WD and AD on spatial frequency and contrasts sensitivity of OKR using a virtual optomotor system (Prusky *et al.*, 2004).

First we investigated the effects of WD on visual acuity. OKR movement thresholds obtained after visual stimulation of either the right or left eye were measured daily for a period of 10 days. Baseline values of WD mice (n=4, **Figure 10a, b**) were always measured before WD, whereas values measured on day 0 represent measurements obtained 4-5 hours after the surgery for WD. Control mice (n=4) remained untreated. Quantitative analysis using one-way ANOVA with repeated measurements revealed significant influences of group (F_1,6_=1165.94, *p<*0.0001) and time (F_11,66_=46.45, p<0.0001) and a significant interaction between the two (F_11,66_=46.45, p<0.0001). Post hoc analysis showed that reflex sensitivity for spatial frequency was unchanged in control animals (n=4) over the whole time period tested. However, in the WD group (n = 4), there was a significant gradual enhancement of spatial frequency sensitivity of the OKR reaching a peak 3 days after WD, about 12% above control level (3 days: control vs WD: 0.40±0.001 (cpd(cycles per degree)) vs 0.45±0.0023 (cpd), *p*<0.0001; unpaired *t-* test followed by Bonferroni correction; **Figure 10a**). Spatial frequency thresholds levels then dropped down over the next two days, but persisted at a level about 5% above control values up to 10 days after WD (10 days: control vs WD: 0.40±0 (cpd) vs 0.42±0.00086 (cpd), *p*<0.0001; unpaired *t*-test followed by Bonferroni correction, **Figure 10a**). Contrast sensitivity of OKR was measured in the same control and WD mice at 0.2 cpd. Group (F_1,6_=656.48, p<0.0001) and time (F_11,66_=69.23, p<0.0001) had a significant influence on contrast thresholds and there was a significant interaction between both (F_11,66_=69.23, p<0.0001, one-way ANOVA with repeated measurements). While contrast thresholds of control mice did not change over the whole time period tested, they gradually increased by almost 100% until day 3 after WD (3 days: control vs WD: 13.03±0.16 vs 26.55±0.74, *p*<0.0001, unpaired t-test followed by Bonferroni correction, **Figure 10b**) suggesting a substantial enhancement of contrast sensitivity due to WD. Subsequently, OKR contrast thresholds decreased again but then remained about 50% above control values between 4 and 10 days after WD (10 days: control vs WD: 13.73±0.26 vs 20.31±0.10; *p*<0.0001, unpaired t-test followed by Bonferroni correction; **Figure 10b**). Taken together, our results suggest that WD cross-modally improves behavioral OKR spatial frequency and contrast sensitivity.

**Figure 10:**
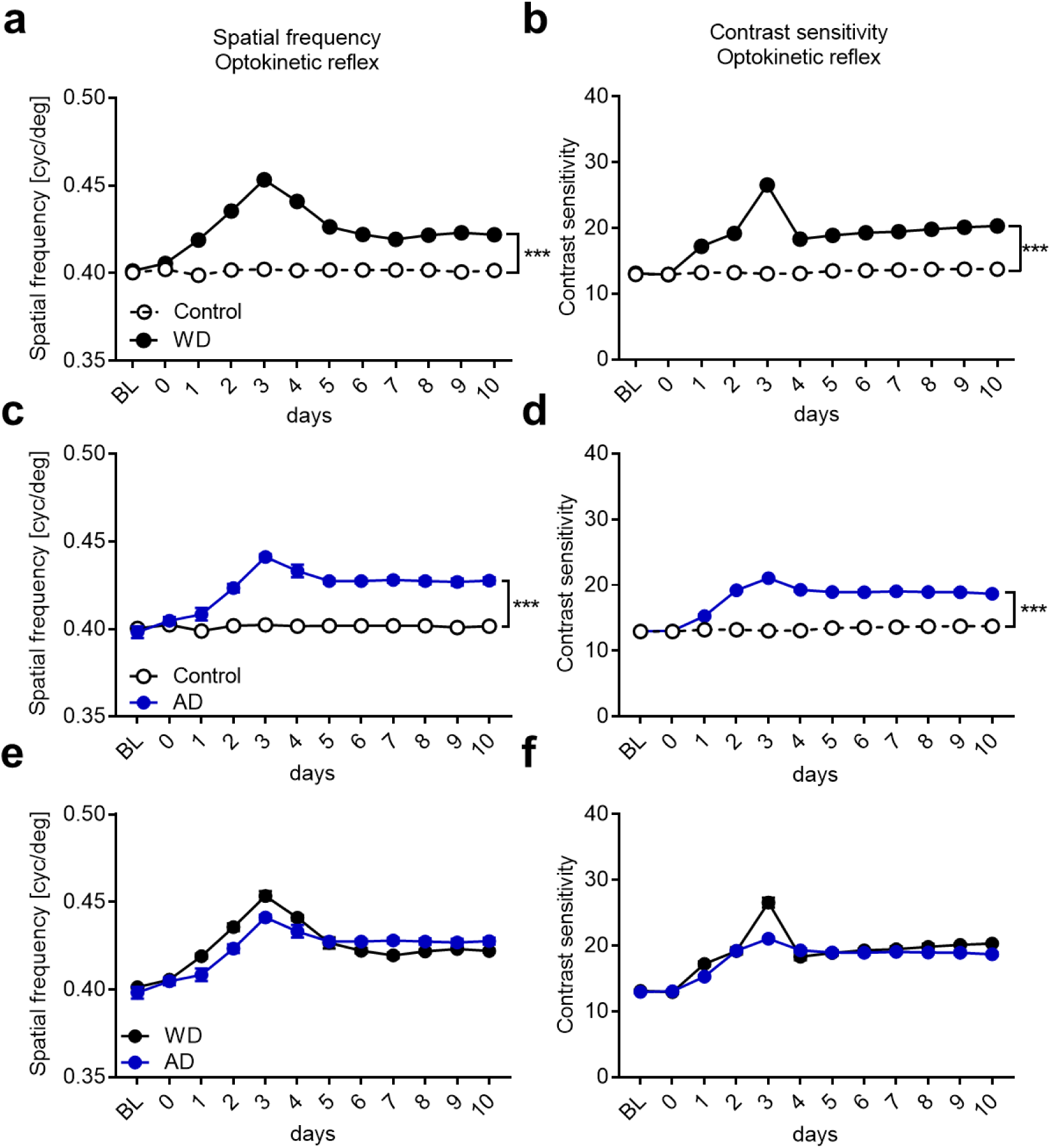
Both WD and AD cross-modally provoke a potentiation of the visual OKR. (**a**) In control mice (n=4), spatial frequency thresholds did not change over the whole time period tested. However, after WD (n=4) there was a marked improvement of the spatial frequency sensitivity which reached a peak on day 3. Subsequently, spatial frequency thresholds levels slightly decreased and remained at a stable level above control values until 10 days after WD. (**b**) Contrast sensitivity of the OKR in control mice remained unchanged over 10 days. After WD, contrast thresholds massively improved until day 3. After a slight decrease at 4 days after WD, values then remained at a stable level above control values until 10 days after WD. (**c**) We used values of the same control mice like in Figure 4a. AD (n=4) led to a marked increase of spatial frequency thresholds peaking at 3 days. Subsequently, spatial frequency sensitivity slightly decreased until day 5 after AD and remained at a stable level above control values until 10 days after AD. (**d**) Contrast thresholds markedly increased until day 3 after AD which was followed by a slight decrease to a stable level above contrast values of control mice. (**e, f**) WD or AD led to similar improvements of the OKR. Open and filled circles represent mean values together with the s.e.m. However, vertical lines of s.e.m. are often occluded by data symbols. ***p<0.001

As a next step, we examined whether AD (n=4) also affects the visual OKR. Quantitative analysis using a one-way ANOVA with repeated measurements showed that group (F_1,6_=654.78, p<0.0001) and time (F_11,66_=24.3, p<0.0001) had a significant influence on spatial frequency thresholds. In addition, we found a significance interaction between group and time (F_11,66_=20.65, p<0.0001, one-way ANOVA with repeated measurements). As shown in **Figure 10c** there was a gradual increase of spatial frequency sensitivity until 3 days after AD (3 days: control vs AD: 0.40 ±0.001 (cpd) vs 0.44±0.002; *p*<0.0001; unpaired *t-*test followed by Bonferroni correction; **Figure 10c**) which was followed by a slight decrease over the next two days to a stable level above control values until day 10 after AD (10 days: control vs AD: 0.40±0 vs 0.43±0.002, *p=*0.0002; unpaired *t-*test followed by Bonferroni correction; **Figure 10c**). Group (F_1,6_=317.79, p<0.0001) and time (F_11,66_=143.39, p<0.0001) also had a significant influence on contrast thresholds and there was a significant interaction between both (F_11,66_=112.92, p<0.0001, one-way ANOVA with repeated measurements). Contrast thresholds also significantly increased until 3 days after AD (3 days: control vs AD: 13.03±0.16 vs 12.06±0.16, *p*<0.0001, unpaired t-test followed by Bonferroni correction; **Figure 10d**) and then remained at a higher level above control measurements until day 10 (10 days: control vs AD: 13.73±0.26 vs 18.7±0.21, *p*<0.0001, unpaired t-test followed by Bonferroni correction; **Figure 10d**). These data indicate that AD can cross-modally improve OKR sensitivity, too. Notably, both spatial frequency and contrast sensitivity changes found after AD were similar to changes observed after WD (**Figure 10e, f**). Taken together, our results strongly suggest that the deprivation of a non-visual modality leads to marked improvements of subcortically mediated visual behavior.

### Cross-modally induced enhancements of the OKR are partially V1 dependent

So far we describe that both WD and AD lead to a potentiation of the OKR. Interestingly, the highest values of OKR thresholds of spatial frequency and contrast as well were obtained 3 days after WD or AD, and thus, exactly at the same time point as the visually driven V1 activity peaks. These results suggest that V1 might be involved in mediating OKR potentiation. In order to address this issue we combined WD and bilateral V1 aspiration (WD+V1aspi, n=3) and measured spatial frequency and contrast thresholds of the OKR over the following 10 days. In mice of the control group we only aspirated V1 bilaterally (V1aspi only, n=3). Baseline values were always measured before V1 aspiration, measurements at day 0 were obtained 4-5 h after WD and V1 aspiration.

For aspiration surgery we located the correct position of V1 using intrinsic signal imaging. **Figure 11a** depicts a representative visually evoked retinotopic polar map of V1, which was merged with a picture of the cortical blood vessel pattern of a normal mouse. **Figure 11b** (left) shows the corresponding amplitude map. We then aspirated V1, guided by blood vessel landmarks, through a small trepanation and performed a second optical imaging session to validate the efficiency of the surgery. As expected, after V1 aspiration visually evoked responses in the V1 area were completely abolished (**Figure 11b**, right, not quantified). These experiments confirm that the surgery for V1 aspiration was efficient and reliable since it completely abolished visually elicited V1 activity. **Figure 11c** shows a representative example of a brain slice 10 days after the aspiration of V1.

**Figure 11:**
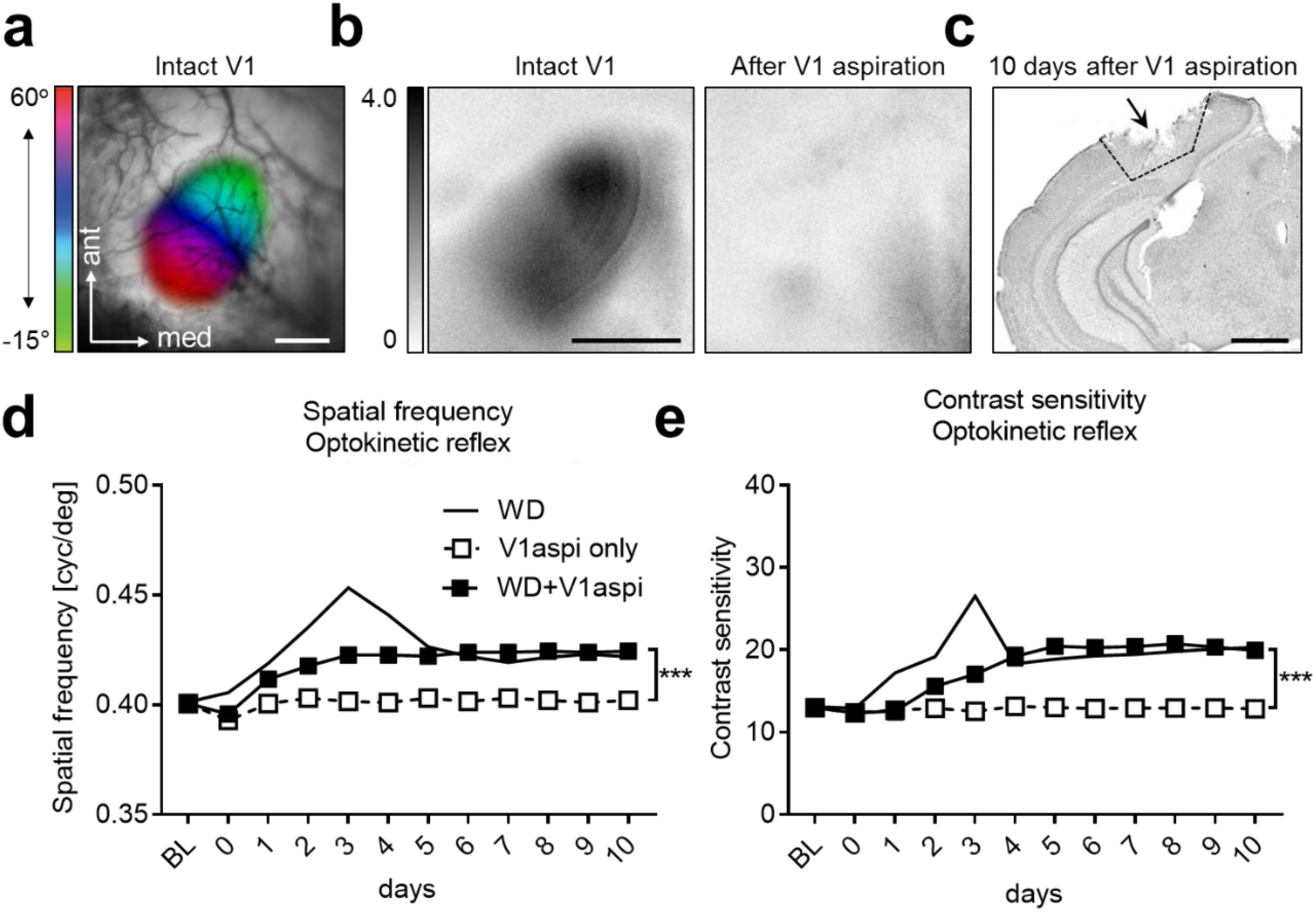
Aspiration of V1 reveals V1 contribution to cross-modally induced enhancements of the OKR. (**a**) V1 was located using intrinsic signal imaging. (**b**) Representative V1 amplitude maps elicited by visual stimulation before and after V1 aspiration. It is clearly visible that after V1 aspiration visually evoked cortical responses were absent demonstrating the efficiency of the aspiration surgery. (**c**) Nissl stained brain slice obtained 10 days after V1 aspiration. (**d**) Spatial frequency thresholds remained unchanged in mice after V1 aspiration only (n=3). After combined WD and V1 aspiration (n=3) spatial frequency sensitivity slightly increased until day 3 and remained at this level for the remaining 7 days. Interestingly, the strong peak of spatial frequency thresholds obtained 3 days after WD only (n=4) was absent in mice after WD and V1 aspiration whereas the long-lasting improvement was almost identical in mice of both groups. (**e**) In WD only, V1aspi only and WD+V1aspi mice, the course of contrast sensitivity thresholds closely followed the course of spatial frequency thresholds in mice of the same groups. Taken together our data suggest that WD leads to V1 dependent and V1 independent improvements of the OKR. Open and filled squares represent mean values together with the s.e.m. However, vertical lines of s.e.m. are often occluded by data symbols.

Quantitative analysis using a one-way ANOVA with repeated measurements showed that group (F_1,4_=77.27, p=0.001) and time (F_11,44_=32.83, p<0.0001) had a significant influence on spatial frequency thresholds. In addition, there was a statistically significant interaction between group and time (F_11,44_=18.00, p<0.0001, one-way ANOVA with repeated measurements). As shown in **Figure 11d** the spatial frequency sensitivity of mice in that we only aspirated V1 (n=3) remained almost unchanged for 10 days. However, if we combined WD and V1 aspiration, spatial frequency thresholds of the OKR slightly increased until day 3 and remained at this level for the whole time period tested (3 days: V1aspi only vs WD+V1aspi: 0.40±0.001 vs 0.42±0.001, 0.004; 10 days: 0.40±0.001 vs 0.42±0.001, *p=*0.002; paired *t-*test followed by Bonferroni correction; **Figure 11d**). Interestingly, the peak of spatial frequency sensitivity at day 3, which we obtained in mice that only received a WD, was abolished in animals after concurrent WD and V1 aspiration. In contrast, the stable level of increased spatial frequency thresholds (between day 5 and 10) which was present in WD mice with combined V1 aspiration was practically identical to the increased stable level reached after WD only. Thus, these results suggest that the marked initial increase and decrease of spatial frequency sensitivity during the first 5 days after WD is mediated by V1. However, the long lasting improvement of spatial frequency thresholds after WD appeared to be V1 independent.

A similar result was obtained for contrast sensitivity of the OKR. Quantitative analysis using a one-way ANOVA with repeated measurements revealed significant influences of group (F_1,4_=44.05, p<0.003) and time (F_11,44_=37.08, p<0.0001) and a significant interaction between both (F_11,44_=31.83, p<0.0001). In mice in which we only aspirated V1 contrasts thresholds remained unchanged for 10 days (**Figure 11e**). When we combined WD and V1 aspiration, contrast thresholds gradually increased until day 5 and remained at this level for the following 5 days (5 days: control vs WD: 13.03±0.79 vs 20.43±0.54, *p=*0.02; 10 days: 12.84±0.67 vs 19.94±0.29, *p=*0.008; paired *t-*test followed by Bonferroni correction; **Figure 11e**). However, the peak of contrast sensitivity found in animals which only received a WD at day 3, was not present in mice after combined WD and V1 aspiration whereas the stable level of enhanced contrast thresholds (4-10 days) was almost identical in animals of both groups (**Figure 11e**). These data suggest that the transient and strong improvement of contrast thresholds of the OKR found in mice after 3 days of WD alone is mediated by activity changes observed in V1. In contrast, the long lasting enhancement of contrast sensitivity does not require V1. Taken together, our results make it reasonable to conclude that WD improves subcortically mediated visual spatial frequency and contrast sensitivity in both a V1 dependent and V1 independent manner.

### Cross-modally induced potentiation of OKR requires NMDA receptor activation

We next examined whether NDMA receptors contribute to improvements of the OKR. For this, WD mice received daily CPP injections (WD+CPP; n=4) and we measured spatial frequency and contrast thresholds again for 10 days. In these mice both spatial frequency and contrast sensitivity remained unchanged over the whole time period tested and were not different from values of untreated control animals (n=4) (spatial frequency: group: F_1,6_=4.84, p=0.07; time: F_11,66_=1.02, p=0.44; interaction: F_11,66_=1.961, p=0.149; contrast sensitivity: group: F_1,6_=1.63, p=0.35; time: F_11,66_=6.8, p=0.008; interaction: F_11,66_=2.097, p=0.16; one-way ANOVA with repeated measurements; **Figure 12a, b**). Hence, CPP administrations abolished both the visual cortex dependent and independent improvements of the OKR thresholds observed in WD mice. These results suggest that potentiation of the OKR induced by the deprivation of a non-visual sense also depends on NMDAR activation, like shown above for V1 activity changes after WD.

**Figure 12:**
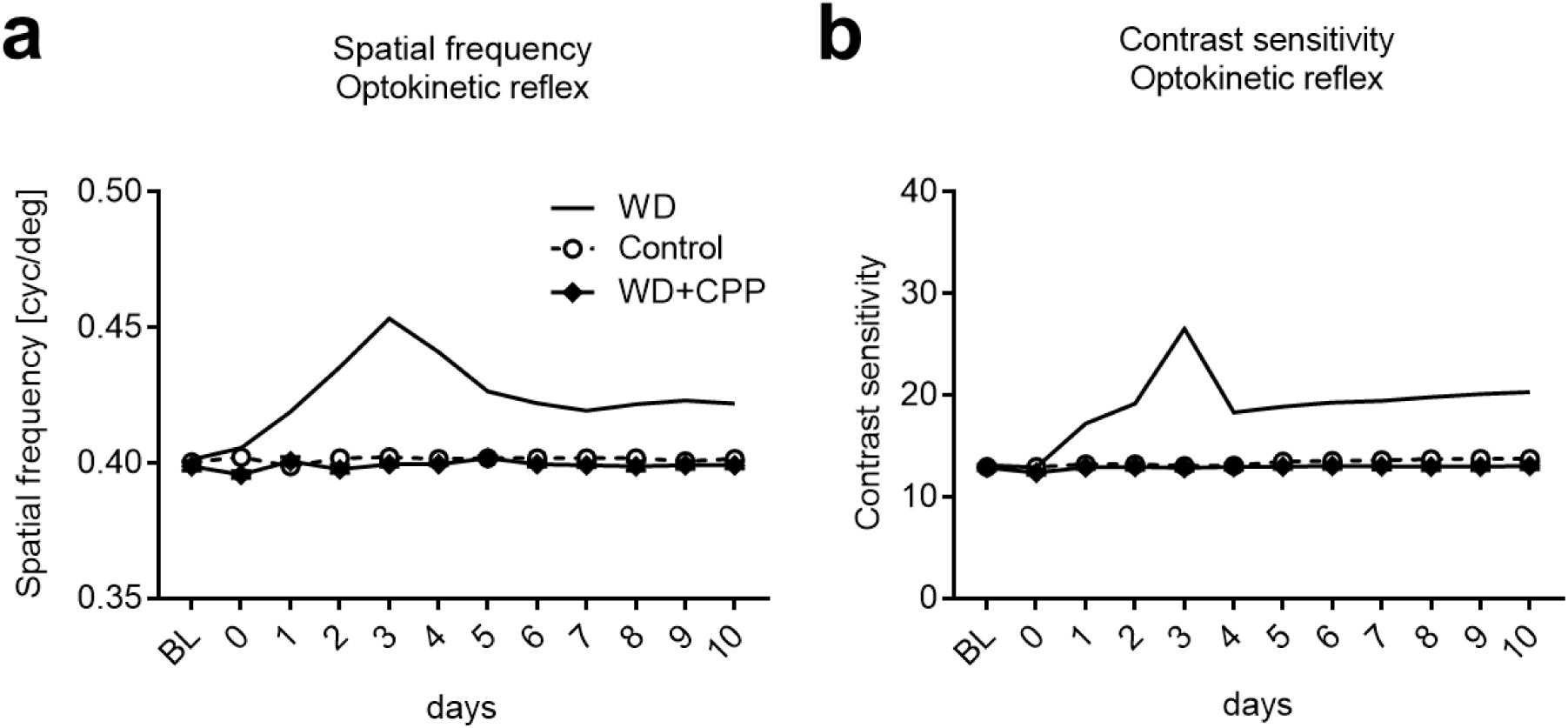
Cross-modally induced OKR potentiation requires NMDAR activation. (**a, b**) In both control mice (n=4) and WD mice which received daily injections of CPP (n=4) spatial frequency and contrast thresholds remained completely unchanged for 10 days. Hence, blocking NMDARs abolished both V1 dependent and V1 independent potentiation of OKR thresholds found after WD only.

## Discussion

In the present study we investigated the effects of WD and AD on visually evoked V1 responses and visually mediated behavior in fully adult mice. We found that both WD and AD transiently shifted the OD in V1 towards the input through the ipsilateral eye. These changes required patterned vision through the ipsilateral eye and were accompanied by an increase of the E/I ratio in V1. Moreover, V1 activity changes partially mediated potentiation of the OKR, a visual behavior predominantly mediated by subcortical structures. These results indicate that the late-onset loss of a non-visual sensory modality dramatically alters neuronal processing in different structures of the visual pathway.

It has been demonstrated that prolonged MD (5-7 days) in “young adult” mice leads to a shift of the OD which is mediated by an increase of V1 activity elicited by open (ipsilateral) eye stimulation (Sawtell *et al.*, 2003; Sato & Stryker, 2008; Ranson *et al.*, 2012). However, this type of “adult” cortical plasticity is completely absent in mice older than 110 days (Lehmann & Lowel, 2008), but can be reinstalled by treatments which change the cortical E/I ratio in favor of excitation (He *et al.*, 2006; Fu *et al.*, 2015). We here show that 3 days of WD or AD in mice older than 110 days lead to V1 activity changes which resemble the type of OD plasticity in younger mice, since OD changes were mediated by an increased V1 responsiveness to ipsilateral eye stimulation (**Figure 3, 4**). This shift in OD was accompanied by an increase of the E/I ratio. By decreasing the E/I ratio with diazepam, the OD shift after WD was abolished. Likewise, after blocking NMDARs, OD plasticity was also abolished (**Figure 9**). Thus, our data indicate that the deprivation of a non-visual modality cross-modally set V1 back into a plastic stage where visual experience, can re-shape V1 circuits. However, in contrast to the classical more than 50 years old paradigm that MD induces alterations of OD (Wiesel & Hubel, 1963; Hubel & Wiesel, 1970; Gordon & Stryker, 1996), the cross-modally induced OD shifts described here took place without visual deprivation. Thus, to the best of our knowledge, we demonstrate here for the first time that, at least in mice, OD can be altered by deprivations of non-visual sensory modalities, too.

Consistent with the finding that OD plasticity in young mice requires NMDAR activation (Sawtell *et al.*, 2003; Sato & Stryker, 2008), we show that this is also the case for cross-modally induced OD changes (**Figure 9**). This result is also in line with our recent study where we demonstrated that cross-modally restored OD plasticity after 7 days of MD also depends on the NMDAR (Teichert *et al.*, 2018b), suggesting that NMDAR activation plays a pivotal role in mediating cross-modal plasticity in spared primary sensory cortices. However, as we administrated the NMDAR blocker CPP systemically, we cannot make statements on the precise location where this receptor is required to mediate cross-modal adaptations. It might be the visual cortex (Sawtell *et al.*, 2003; Sato & Stryker, 2008) but, since a recent study could demonstrate that OD shifts already take place in in the lateral geniculate nucleus (Jaepel *et al.*, 2017), it is possible that NMDARs are already required in early structures of the visual pathway.

Previous studies have demonstrated that the deprivation of one sense for only a few days strengthens thalamo-cortical and layer 4 to 2/3 synapses in a spared primary sensory cortex (Jitsuki *et al.*, 2011; Petrus *et al.*, 2014; Petrus *et al.*, 2015). In accordance with this finding, we here demonstrate that WD strengthens V1 layer 4 synapses (**Figure 7**). At least a part of these strengthened synapses most likely represent thalamo-cortical synapses for several reasons: First, strengthening of thalamo-cortical synapses leads to increased sensory driven responsiveness of primary sensory cortices (Heynen & Bear, 2001; Petrus *et al.*, 2014). This is in line with our imaging results, as evoked V1 activity was increased 3 d after WD. Second, visual input is required to mediate V1 activity changes (**Figure 6**) suggesting an involvement of synapses on the visual pathway. And third, NMDA receptors are involved in mediating this cross-modal effect (**Figure 9**). Together, these results also suggest that cross-modal plasticity in the spared sensory cortex is a form of experience dependent synaptic plasticity similar to long-term potentiation (LTP) (Lee & Whitt, 2015). Furthermore, previous studies reported that 7 days of sensory deprivation lead to a decrease of AMPAR mediated miniature EPSC amplitudes in layers 2/3 of the spared primary sensory cortex (Goel *et al.*, 2006; He *et al.*, 2012). Hence, it was speculated that an initial strengthening of synapses in the remaining sensory cortices (after 2-3 days) is followed by a decrease in synaptic transmission after 7 days of sensory deprivation (Jitsuki *et al.*, 2011). Our functional data of V1 responsiveness support this hypothesis as 7 days after WD or AD V1 responses were completely restored back to baseline levels (**Figure 3, 4)**. Thus, our data suggest that the restoration of normal OD levels is mediated by homeostatic mechanisms like synaptic down-scaling (Goel *et al.*, 2006; He *et al.*, 2012) and/or cross-modally induced reduction of lateral input strength in layers 2/3 as shown previously (Petrus *et al.*, 2015).

We could recently show that 7-12 days of WD dramatically improved V1 mediated visual acuity and contrast sensitivity, as measured by behavioral visual water task experiments (Teichert *et al.*, 2018a). In the present study we demonstrate that both WD and AD also provoked a fast and a long-lasting improvement of the so called optokinetic reflex (OKR), another type of visual behavior mainly mediated by subcortical structures including the cerebellum and vestibular nuclei (Liu *et al.*, 2016). Previous studies demonstrated enhanced OKR sensitivity in rodents after MD (Prusky *et al.*, 2006), vestibular impairments (McCall & Yates, 2011; Liu *et al.*, 2016) or daily threshold testing from eye opening into adulthood (Prusky *et al.*, 2008). Here, however, we provide the first evidence that OKR improvements can also be induced by depriving somatosensation or hearing (**Figure 10, 11**). We found that both, WD and AD resulted in three distinct phases of altered OKR sensitivity: During the first phase, OKR sensitivity markedly increased and peaked at 3 days after WD or. The second phase was characterized by a drop of OKR thresholds, lasting for 2-3 days, but still remained above control levels. Phase one and two appeared to be V1 dependent. These result are in line with previous studies demonstrating that V1 is involved in enhancements of OKR sensitivity (Prusky *et al.*, 2006; Prusky *et al.*, 2008; Liu *et al.*, 2016). Interestingly, the transient activity peak in V1 3 days after WD or AD temporally matched the peak of OKR improvements. These results suggest that cortico-fugal projections might transmit cross-modally induced V1 activity changes to subcortical structures which then act to mediate the compensatory potentiation of OKR. This hypothesis is supported by the findings that cortico-fugal projections can indeed modulate sensory induced behaviors (Liang *et al.*, 2015; Xiong *et al.*, 2015). However, the third phase, where OKR sensitivity remained at a stable level above baseline for at least 10 days, was V1 independent as mice with removed whiskers (WD) and aspirated V1 also reached this enhanced level (**Figure 11**). Together with our previous finding that WD leads to an enhancement of visual performance in the visual water task (Teichert *et al.*, 2018a), these data indicate that the deprivation of non-visual senses provokes a general long-lasting compensatory improvement of visually mediated behaviors. Interestingly, like cross-modally provoked V1 activity changes, potentiation of the OKR could be completely abolished by antagonizing NMDARs (**Figure 12**). These results then suggest that NMDAR in different structures of the visual pathway are instrumental in the mediation of cross-modal effects.

In summary, we could demonstrate that the deprivation of non-visual sensory modalities transiently changes OD in V1. We postulate that reducing either somatosensory or auditory input cross-modally re-installs V1 plasticity in fully adult mice, allowing visual inputs to compensatorily re-shape V1 circuits. While further studies are needed to clarify the precise mechanisms underlying this novel and surprising finding, the present results already emphasize the power of cross-modal plasticity to re-open a window of high plasticity in the fully adult cortex far beyond any sensory critical period.

## Acknowledgements

Thanks are due to PD Dr. Konrad Lehmann for his help with statistical analysis, Elisabeth Meier for excellent technical assistance, Tanja Herrmann for her help with electrophysiological recordings and Sandra Eisenberg for animal care.

## Author contribution statement

M.T., M.I. and J.B. designed the study

M.T., M.I., L.L., C.A.H., F.W., C.W. performed experiments

M.T.; M.I., L.L., JB analyzed data

M.T. and J.B. wrote the paper

